# Apoplastic class III peroxidases PRX62 and PRX69 regulate ROS-homeostasis and cell wall associated extensins linked to root hair growth at low-temperature in *Arabidopsis thaliana*

**DOI:** 10.1101/2021.08.20.456256

**Authors:** Javier Martinez Pacheco, Philippe Ranocha, Luciana Kasulin, Corina M. Fusari, Lucas Servi, Lucía Ferrero, Victoria Berdion Gabarain, Juan Manuel Peralta, Cecilia Borassi, Eliana Marzol, Diana Rosa Rodríguez-Garcia, Yossmayer del Carmen Rondón Guerrero, Mariana Carignani Sardoy, Javier Botto, Claudio Meneses, Federico Ariel, Ezequiel Petrillo, Christophe Dunand, José M. Estevez

## Abstract

Root Hairs (RHs) growth is highly influenced by endogenous as well as by external environmental signals that coordinately regulate its final cell size. RHs actively expand the root surface responsible for nutrient uptake and water absorption. We have recently determined that RH growth was unexpectedly boosted when *Arabidopsis thaliana* seedlings are cultivated at low temperatures. It was proposed that RH growth plasticity in response to low temperature was linked to a reduced nutrient availability in the media. Here, we explored the molecular basis of this strong RH growth response by using the Genome Wide Association Studies (GWAS) approach on *Arabidopsis thaliana* natural accessions. We identified the poorly characterized PEROXIDASE 62 (PRX62) as a key protein triggering this conditional growth under a moderate low-temperature stress. In addition, we identified the related protein PRX69 as an important factor in this developmental process. The *prx62 prx69* double mutant and the *PRX62* and *PRX69* over-expressing lines showed contrasting RH phenotypes, peroxidase activities and cyt/apoReactive Oxygen Species (ROS) levels. Strikingly, a cell wall protein extensin (EXT) reporter revealed the effect of peroxidase activity on the EXT cell wall association at 10°C in the RH apical zone. EXT cell wall insolubilization was enhanced at 10°C, which was completely abolished under the PRX inhibitor salicylhydroxamic acid (SHAM) treatment. Finally, we demonstrated that the *Root Hair defective 6-like 4* (RSL4) bHLH family transcription factor directly controls the expression of PRX69. Collectively, our results indicate that both PRX62 and PRX69 are key apoplastic PRXs that modulate ROS-homeostasis and cell wall EXT-insolubilization linked to RH elongation at low-temperature.

## INTRODUCTION

Root hairs (RH) have emerged as an excellent model system for studying cell size regulation since they can elongate several hundred-fold their original dimensions. The rate at which cells grow is determined both by cell-intrinsic factors as well as by external environment signals. RHs represent an important proportion of the surface root area, crucial for nutrient uptake and water absorption. RH growth is controlled by the interaction of several proteins, including the bHLH transcription factor (TF) RSL4 (*Root Hair Defective Six-like 4*), which defines the final RH length (Datta et al. 2015; Mangano et al. 2017) as well as the related TF RSL2 (*Root Hair Defective 6 Six-like 2*; Bhosale et al. 2018; Mangano et al. 2018). Together with the developmental and genetic pathways, several hormones are important modulators of RH cell growth (Lee & Cho 2013; Velasquez et al. 2016; Zhang et al 2016; Mangano et al. 2017). In addition, abnormal Reactive Oxygen Species (ROS) accumulation in RHs triggers either exacerbated growth or cell bursting. Exogenous H_2_O_2_ inhibited RH polar expansion, while treatment with ROS scavengers (e.g., ascorbic acid) caused RH bursting, reinforcing the notion that a balanced ROS-homeostasis is required to modulate cell elongation by affecting cell wall properties. Accordingly, apoplastic ROS (_apo_ROS) produced in the apoplast (specifically _apo_H_2_O_2_) coupled to apoplastic Class III peroxidase (PRX) activity directly affect the degree of cell wall crosslinking (Passardi et al. 2004) by oxidizing cell wall compounds and leading to the stiffening of the wall in peroxidative cycles (PC) (Orman-Ligeza et al. 2016). In addition, _apo_ROS coupled to PRX activity enhances non-enzymatic wall loosening by producing oxygen radical species (e.g., ^●^OH) and promoting polar-growth in hydroxylic cycles (HC) (Dunand et al. 2007). Finally, PRXs also contribute to the production of superoxide radical (O_2_^●-^) pool together with NADPH oxidase/respiratory burst oxidase homolog (RBOH) proteins by oxidizing singlet oxygen in the oxidative cycle (OC), thereby affecting _apo_H_2_O_2_ levels. Given their multiple enzymatic activities *in vivo,* apoplastic PRXs emerge as versatile regulators of rapid cell elongation. Assigning specific functions to each of the numerous PRXs (73 encoded in Arabidopsis; Valerio et al. 2004; and even more in other plant types, e.g. 138 encoded in Rice; Passardi et al. 2004a) has been challenging. Recently, three PRXs possibly linked to Tyr-crosslinking of cell wall extensins (EXTs), PRX01, PRX44 and PRX73, were characterized as important regulators of RH growth under low-nutrient conditions (Marzol et al. 2021). These RH specific PRXs are under the direct control of the TF RSL4, a master regulator of RH cell size (Yi et al. 2010; Datta et al. 2015; Mangano et al. 2017). In addition, other PRXs were postulated to crosslink EXTs in aerial plant tissues. PRX09 and PRX40 were proposed to crosslink EXTs during tapetum development, and both, PRXs were able to crosslink EXT23 in transient expression experiments (Jacobowitz et al. 2019).

Although there is a fairly well-known mechanistic view of how RH cell expands, the environmental signals that trigger the cell elongation process remain currently unknown. Due to its important role in root physiology, it has been anticipated that RH would be highly susceptible to environmental stresses such as heat or moderate temperature increase, which trigger extensive DNA methylation, transcriptomic and proteomic changes (Valdés-López et al. 2016; Quint et al. 2016; Hossain et al. 2017). Although RH development during cold acclimation remains largely unexplored, it has been observed that many RH-related genes respond to cold in the whole plant or seedlings (Maruyama *et al*. 2004; Hannah, Heyer & Hincha 2005; Barah et al. 2013). It is known that plants may perceive cold by a putative receptor at the cell membrane and initiate a signal to activate the cold-responsive genes and transcription factors for mediating stress tolerance (Thomashow 1999; Penfield 2008; Ding *et al*., 2019; Nurhasanah Ritonga and Chen, 2020; Leuendorf *et al*., 2020). Previously, we have shown that the plant long noncoding RNA (lncRNA) *AUXIN REGULATED PROMOTER LOOP* (*APOLO*) recognizes the locus encoding the RH (RH) master regulator RHD6 (*Root Hair Defective 6*) and controls *RHD6* transcriptional activity leading to cold-enhanced RH elongation through the consequent activation of *RSL4* (Moison et al. 2021). In addition, *APOLO* is able to bind and positively control the expression of several cell wall EXTENSIN (EXT) encoding genes, including *EXT3*, a key regulator for RH growth (Martinez-Pacheco et al. 2021). Unexpectedly, our previous results indicate that the low-temperatures (10°C) are able to trigger an exacerbated RH growth compared with cell expansion at room temperature (Moison et al. 2021; Martinez-Pacheco et al. 2021). To explore the molecular basis of this strong growth response, we conducted Genome Wide Association Studies (GWAS) on *Arabidopsis thaliana* natural accessions and identified the uncharacterized PEROXIDASE 62 (PRX62) as a key protein that regulates the conditional growth under a moderate low temperature stress. In addition, we also identified a second PRX, i.e. PRX69, as an important player in this developmental response. Both, PRX62 and PRX69 are key enzymes to trigger RH growth, likely by participating in a ROS-mediated mechanism of polar cell growth at low-temperatures. The expression of both PRX encoding genes could be under the regulation of RSL4, which has a direct binding to *PRX69* promoter specific regions. Transcriptomic analyses revealed that upon *PRX62* and *PRX69* knockout, several other PRXs and cell wall EXTs encoding genes were differentially expressed, hinting at a compensatory mechanism.

## RESULTS AND DISCUSSION

### PRX62 and PRX69 emerged as positive regulators of RH growth at low-temperatures

In order to identify the natural genetic components involved in RH growth under low-temperature conditions (at 10°C), we analyzed natural *A. thaliana* accessions originated from contrasting environments (Europe, Asia, Africa and North America, **Figure S1**). We assessed RH growth for each seedling accession grown under 22°C for 5 days, and then transferred them to 10°C for 3 days. RH length was the phenotypic trait recorded for each accession, and compared to seedlings grown at 22°C for 8 days, taken as a control. We observed 15-folds range of natural variation for average RH length (148-2218 µm) in the accessions grown at 10°C (**Figure S2A**; **Table S1**) in contrast with a lower variability (∼7-folds) and significantly shorter overall RH cells when seedlings were grown at 22°C (136-1034 µm). There is a strong correlation (R^2^=0.981) for RH length from accessions grown at 22°C→10°C compared to plants growth at 22°C (**Figure S2B**), indicating that accessions respond in the same manner to a temperature decrease but varying in intensity. Only the most contrasting accession are shown as examples (**Figure 1A**). Thus, moderate low-temperature triggers RH polar-growth across Arabidopsis ecotypes by a yet unknown molecular mechanism. To identify candidate genes involved in RH growth response at moderate low-temperature, we performed a GWAS (GWAPP web tool, Seren et al, 2012) using as input data the RH length recorded only at 22°C or at 22°C →10°C for each accession (**Table S1**). When GWAS was performed measuring RH length obtained at 22°C, no significant associations were identified (**Figure 1B**). On the other hand, after filtering SNPs for a 10% minor allele frequency in the 22°C→10°C RH length GWAS, a leader SNP m190905 (TAIR10 position 15847854) was significantly associated with RH length (LOD [for log of the odds] =6.01, FDR=0.06) with RH length. This SNP is located in the intron of *PEROXIDASE62* (*PRX62*, AT5G39580). Three additional SNPs located in *PRX62* exons, in high linkage disequilibrium with m190905 (r^2^ > 0.7, p<0.001), showed relatively high LOD score of association (m190904/15847644, m190907/15848071, m190909/15848704, LOD ∼ 4.99-4.24, **Figure 1C**). These four SNPs formed seven haplotypes, with two major allele-opposite haplotypes (CTGT, n=79; TGAA, n=18), two haplotypes with very low frequency (CTGA, n=5; TGGA, n=5) and three unique haplotypes (**Figure S3A**). Analysis of variance between the average trait values for all non-unique haplotypes showed that RH length varies among them, having the first and second most frequent haplotypes significantly different values for RH length (**Figure S3B**). In addition, *PRX62* presents two splice variants, differing in the sequence length of the last exon. We then analyzed if these two *PRX62* isoforms can be equally detected and if low-temperature treatment induces a differential expression of any of them. According to **Figure 1D**, only the full-length transcript of *PRX62* (AT5G39580.1) is detectable in the Col-0 that increased up to 2.54 log_2_FC in roots under low-temperature (RNA-seq). This was further confirmed by RT-qPCR (**Figure S4**). Altogether, our results hinted at *PRX62* as a potential key factor in the regulation of RH growth under low-temperature. Interestingly, according to publicly available datasets of whole seedlings (Schlaen et al. 2015), six PRX genes appeared as induced at 10°C; notably *PRX62* and *PEROXIDASE69* (PRX69, AT5G64100) were predicted to be highly expressed in RHs (**Table S2**). *PRX69* also has two predicted variants, the full length AT5G64100.1 and a shorter one AT5G64100.2. By RNA-seq we also confirmed that only the full-length variant of PRX69 is the most expressed one with an small upregulation (by 0.21 log_2_FC) by low temperature although with similar overall transcriptional levels to *PRX62* (**Figure 2D**). This was also confirmed by RT-qPCR (**Figure S4**). Therefore, we decided to characterize in depth both PRXs, PRX62 and PRX69 and their roles in RH growth at 10°C.

**Figure 1.**
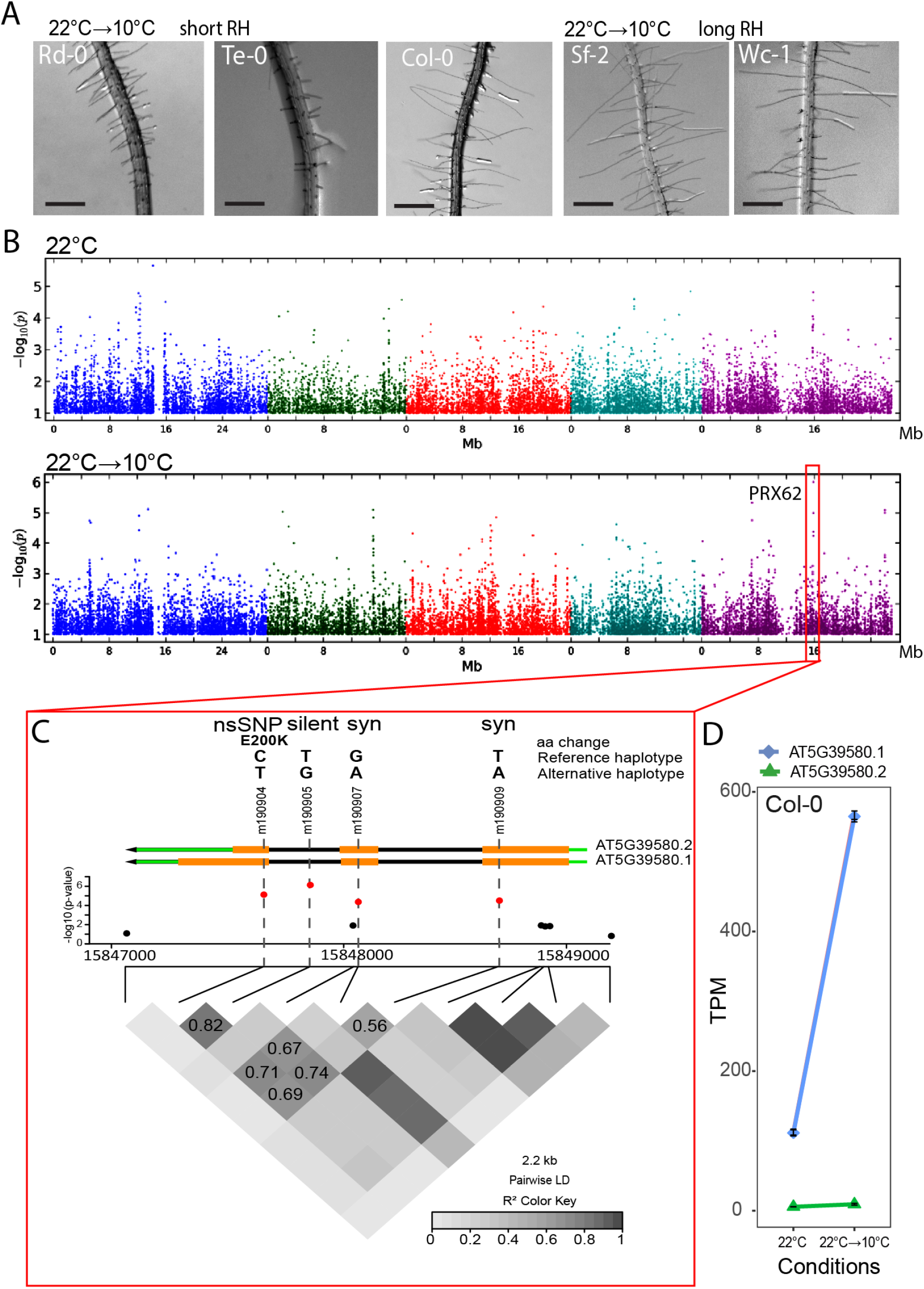
*PRX62* associates with enhanced RH growth under low-temperature condition. (A) Representative accessions of *A. thaliana* showing short (Rd-0 and Te-0) and long root hair (RH) phenotypes (Col-0, Sf-2 and Wc-1) when grown at low temperature (10°C). (B) Manhattan plots for RH length at 22°C (top plot) and at 10°C (bottom plot). Coarse analysis was performed using GWAPP (https://gwapp.gmi.oeaw.ac.at/). Arabidopsis chromosomes are depicted in different colors. The red box in the bottom plot indicates the genomic region significantly associated with root hair length at 10°C. (C) Zoomed-in of the genomic region red-boxed in (**B**). The lead SNP (m190905) and three additional SNPs highly associated with RH length localize within *PRX62* (AT5G39580). *PRX62* splice variants (AT5G39580.1, AT5G39580.2) are depicted in green-orange-black arrows. The four associated SNPs (in red) are in high linkage disequilibrium (LD) with each other, and they are combined into two major and opposite haplotypes in the population (CTGT and TGAA). LD plot is shown as heat-map at the bottom. Haplotypes and type of mutation for these SNPs are indicated at the top of the figure. The SNP m190904 is a non-synonymous SNP for AT5G39580.2 causing a change from Glutamic Acid to Lysine at position 200 (E200K) in the amino acid sequence. (D) The full-length variant of PRX62 (AT5G39580.1) is upregulated at low-temperature (10°C) while the shorter variant (AT5G39580.2) is almost not detected. Expression measured by RNA-seq of *PRX62*. TPM = Transcripts Per Kilobase Million.

**Figure 2.**
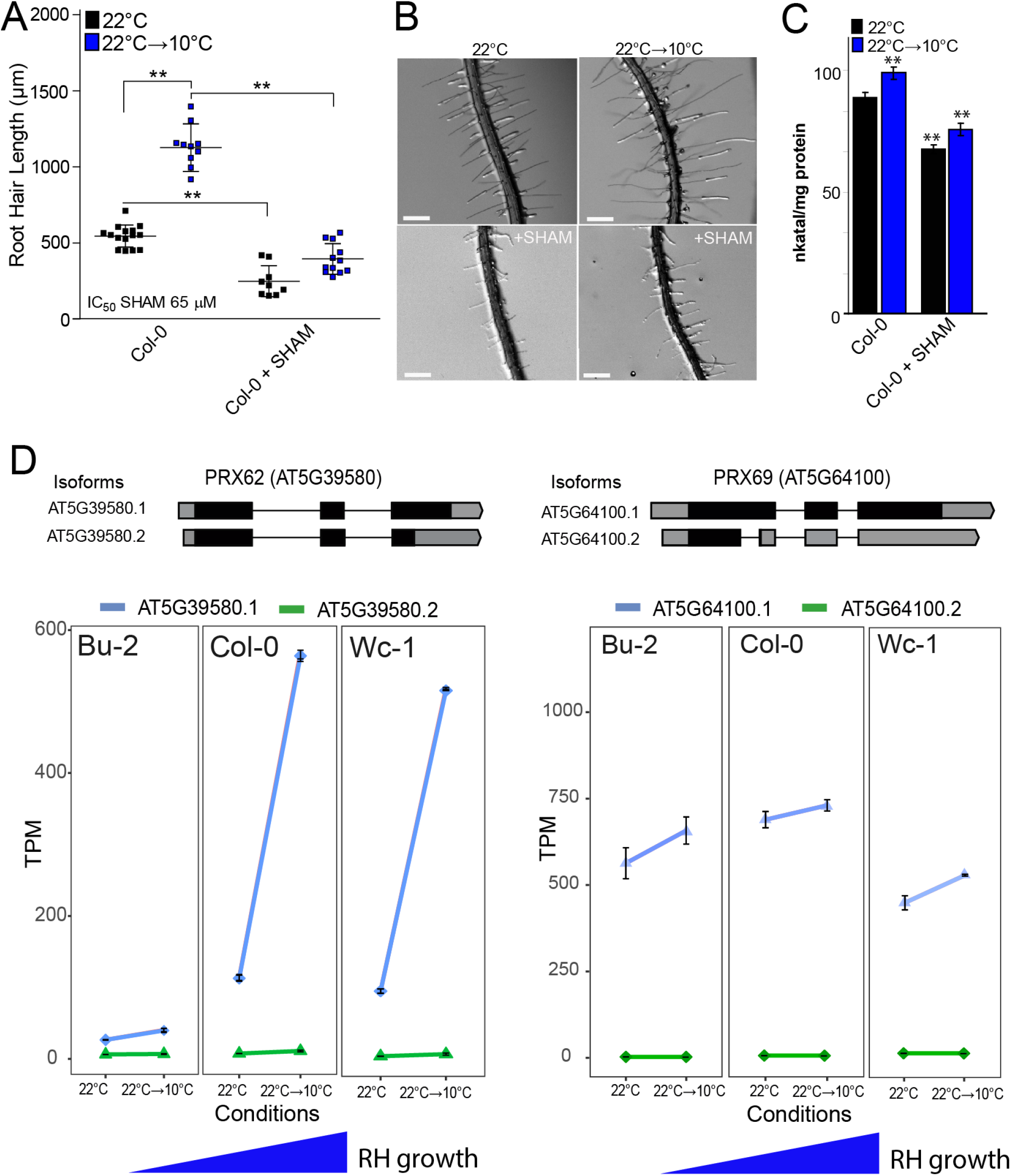
Low-temperature enhanced RH growth requires peroxidase activity and upregulates *PEROXIDASE 62 (PRX62)* expression. (A) RH length phenotype of Col-0 at 22°C or 10°C, with or without the addition of the PRX inhibitor SHAM. Inhibitory Concentration 50 (IC_50_) of RH grown at 22°C was used (65 µM). RH length values are the mean of three replicates ± SD. P-value of one-way ANOVA, (**) P<0.01. (B) Representative images of RH phenotype of Col-0 quantified in (**A**). Scale bars= 0.5 mm. (C) Total root peroxidase activity. Peroxidase activity was assayed using guaiacol/hydrogen peroxide as substrate in root tissues grown with or without 65 µM SHAM, either for 5 days at 22°C or for 5 days at 22°C plus 3 days at 10°C. Enzyme activity values (expressed as nkatal/mg protein) are the mean of three replicates ± SD. P-value of one-way ANNOVA, (**) P<0.01. (D) In contrast to *PRX69*, *PRX62* is differentially expressed at low temperature (10°C) in *Arabidopsis* accessions with contrasting RH phenotypes. Expression measured by RNA-seq of *PRX62* and *PRX69* in three contrasting *Arabidopsis* accessions based on the RH phenotype (short RH in Bu-2 and extra-long RH in Col-0 and Wc-1) detected at 10°C. Isoforms’ schemes were adapted from boxify (https://boxify.boku.ac.at/). TPM = Transcripts Per Kilobase Million.

In agreement with GWAS results, low-temperature-mediated growth requires peroxidase activity since the treatment with salicylhydroxamic acid (SHAM), a peroxidase inhibitor (Kim et al. 2012; Kwon et al. 2015) at inhibitory concentration 50% (IC_50_=65µM) at 22°C was able to repress up to 90% of this growth response at low temperature (**Figure 2A-B**). Accordingly, peroxidase activity in whole roots was significantly lower under the SHAM treatments at both temperatures (**Figure 2C**). We then tested if *PRX62* and *PRX69* expression levels were different between contrasting accessions based on the RH phenotype at 10°C (**Figure 2D**; **Figure S4**). Transcript levels of *PRX62* (after 3 days at 10°C) were positively correlated with the RH length of the given accession, i.e. that the higher the expression of PRX62 at 10°C, the longer the RHs. This implies that high levels of PRX62 in Wc-1 and very low levels in Bu-2 accessions might be linked to the differential RH phenotype detected at low-temperature and suggests that the causal variation for RH length is dependent on PRX62 higher expression (**Figure 2D**; **Figure S4**). On the contrary, *PRX69* transcript levels are higher at 10°C, but they did not show any significant variation across accessions. Altogether, these results suggest that upregulation of *PRX62* transcript levels together with high levels of *PRX69* play an important role in RH growth at low-temperature.

### PRX62 and PRX69 regulate RH growth under low-temperature

The *in silico* analysis of *PRX62* and *PRX69* expression using Tissue Specific Root eFP (http://bar.utoronto.ca/eplant/) showed that both PRX encoding genes were confined to differentiated RH cells with expression in the elongation phase at similar levels than the RH marker *EXPANSIN 7* (**Figure 3A**). Accordingly, the corresponding reporter lines of *PRX62_pro_GFP* as well as *PRX69_pro_GFP* showed high levels of signal in RH cells when grown at 10°C while lower expression was detected at 22°C (**Figure 3B**). When *PRX62* and *PRX69* tagged constructs (*35S_pro_PRX62-TagRFP* and *35S_pro_PRX69-TagRFP*) are transiently coexpressed in *Nicotiana benthamiana* leaves with a plasma membrane marker, both PRXs showed an apoplastic localization (**Figure S5**). Overall, these results confirm that PRX62 and PRX69 are both cold-responsive specific RH class III PRXs that are secreted to the apoplastic space in the cell wall. In order to test if the absence of PRX62 and PRX69 is able to modify growth response at 10°C, we assessed two T-DNA mutants for PRX62 in the Col-0 background (*prx62-1* and *prx62-2*), being a knock-out (*prx62-1*) and a knock-down (*prx62-2*) allele, respectively (Jemmat et al. 2020). In addition, we also characterized two previously reported T-DNA mutants for PRX69 (*prx69-1* and *prx69-2*) (Jemmat et al. 2020). By RNA-seq, we confirm they were absence of transcripts for both PRX62 and PRX69 in these mutants (**Figure S6A**). Only in *prx69-1* when grown at 10°C we found a truncated transcript of PRX69 (**Figure S6B**). The RH phenotype in both *prx62* and prx69 single mutants were similar to Col-0 at 22°C and at 10°C (**Figure 3C**) while the double mutant *prx62-1 prx69-1* showed significantly shorter RHs than Col-0 and any of the single mutants *prx62* and *prx69* at 10°C. The double *prx62-1 prx69-1* mutant showed no detectable transcript levels of both PRXs (**Figure S6C**). The overall peroxidase activity was also partially impaired in single mutants *prx62-1* and *prx69-1* and double mutant *prx62-1 prx69-1* at both growth temperatures, 22°C and 10°C (**Figure 3D**). We then tested RH growth complementation of the *prx62-1 prx69-1* double mutant by expressing either PRX62 or PRX69 coding sequences under 35S promoter (35S_pro_PRX62, 35S_pro_PRX69). The RH growth was restored comparable to Col-0 levels at 10°C only for PRX62 but not for PRX69 (**Figure S7**). This suggests that high levels of *PRX62* but not of *PRX69* are able to compensate the absence of both PRXs in *prx62 prx69* double mutant. To determine whether higher expression of the *PRX62* and *PRX69*-encoding genes are sufficient to trigger changes in RH cell length, we generated a constitutive *35S_pro_PRX62* overexpression lines in the Col-0 background that expressed up to 13-52 folds of transcripts levels of PRX62 as well as the corresponding *35S_pro_PRX69* overexpression lines with 9-11 folds (**Figure S6C**). As expected, PRX62 overexpression resulted in significantly longer RH cells than their respective Col-0 while PRX69 overexpression failed to trigger enhanced growth (**Figure 3E**). This may indicate that PRX62 and PRX69 do not have equal functions in RH growth although both PRXs are required for this enhanced low-temperature growth process. Taken together, these results indicate that the amount of PRX62 and PRX69 proteins linked to their peroxidase activities control RH growth at 10°C.

**Figure 3.**
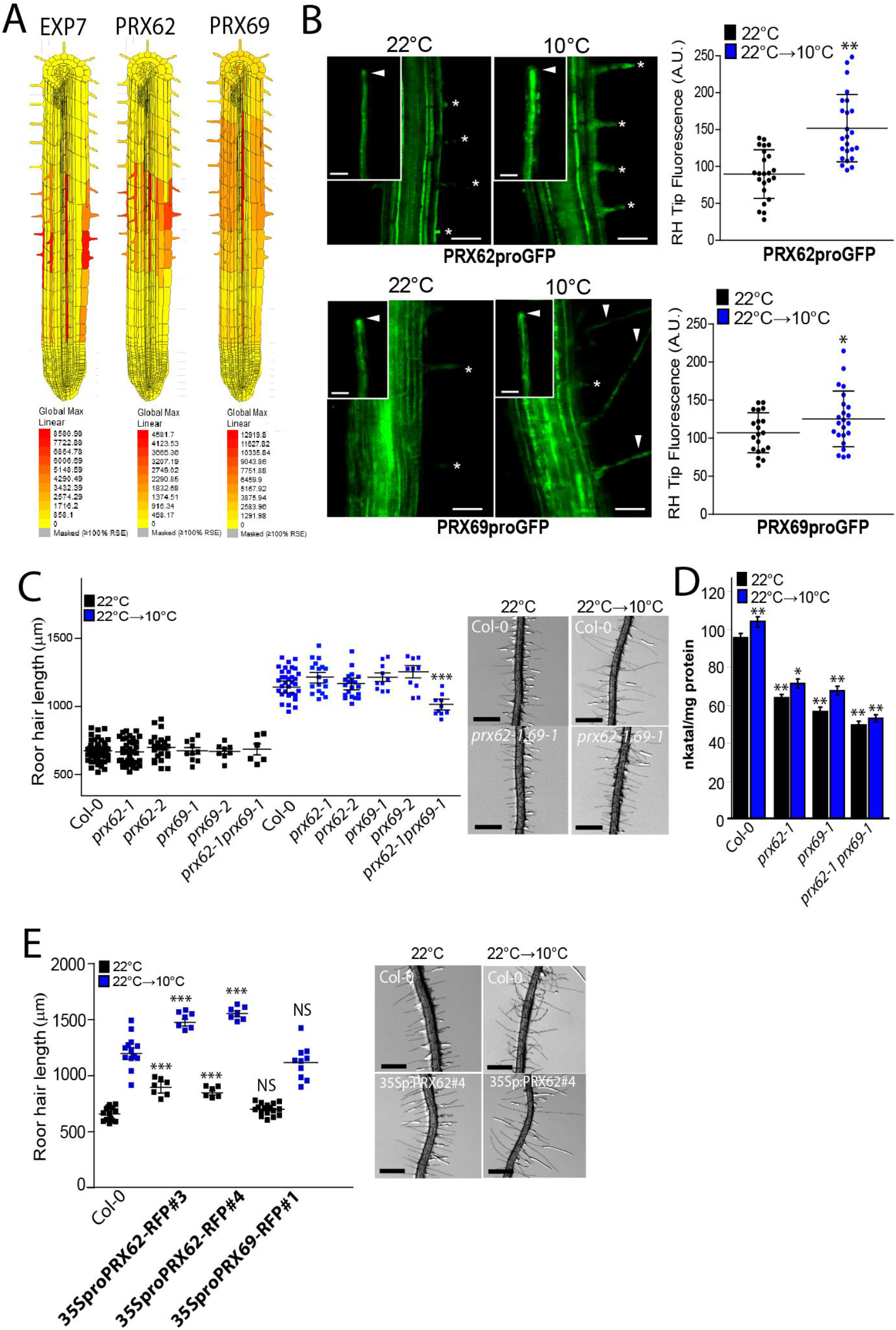
*PEROXIDASE 62* (*PRX62*) and *PEROXIDASE 69* (*PRX69*) regulate RH growth and peroxidase activity under low-temperature conditions. (A) The *in silico* analysis of *PRX62* and *PRX69* gene expression using Tissue Specific Root eFP (http://bar.utoronto.ca/eplant/). The RH marker *EXPANSIN 7* was included for comparison. (B) Transcriptional reporters of *PRX62* (*PRX62_pro_GFP*) and *PRX69* (*PRX69_pro_GFP*) in the root elongation zone and specifically in RHs (RH) grown at 22°C or 10°C. Scale bar = 200 μm. Growing RHs are indicated with asterisks while already grown RHs with arrowheads. On the right, GFP signal is quantified. Fluorescence AU were reported as the mean of three replicates ± SD. P-value of one-way ANOVA, (**) P<0.01, (*) P<0.05. (C) Scatter-plot of RH length of Col-0, *PRX62* mutants (*prx62-1* and *prx62-2*) and *PRX69* mutants (*prx69-1* and *prx69-2*) and double mutant *prx62-1 prx69-1* grown at 22°C or at 10°C. RH length values are the mean of three replicates ± SD. P-value of one-way ANOVA, (***) P<0.001. (D) Peroxidase activity was assayed using guaiacol/hydrogen peroxide as substrate in root tissues from Col-0, *prx62-1* and *prx69-1* seedlings grown either at 22°C or 10°C. Enzyme activity values (expressed as nkatal/mg protein) are the mean of three replicates ± SD. P-value of one-way ANOVA, (**) P<0.001, (*) P<0.05. (E) Scatter-plot of RH length of Col-0, *35S_pro_PRX62*/Col-0 and *35S_pro_PRX69*/Col-0 lines. RH length values are the mean of three replicates ± SD. P-value of one-way ANOVA, (***) P<0.001. NS= non-significant differences.

### The absence of PRX62 and PRX69 induced a deregulation of several PRXs and cell wall EXTs at low-temperature

To better understand the transcriptional changes produced at low temperature in a PRX62- and PRX69-dependent manner, we performed an RNA-seq analysis comparing Col-0 and the double *prx62-1 prx69-1* mutant at 10°C or 22°C. We found a central core of 1544 differentially expressed genes (DEG) at low-temperature grouped into 10 clusters that were misregulated in the double *prx62-1 prx69-1* mutant compared to Col-0 (**Figure 4A**). 1022 genes were upregulated (clusters 1-6) and 522 were downregulated (clusters 7-10) in Col-0 compared to the double *prx62 prx69* mutant. We focused on the largest clusters 1, 2 and cluster 4 (comprised by 873 genes) where the genes upregulated in Col-0 were deregulated in double *prx62 prx69* mutant in response to cold. In these gene clusters, overrepresented GO terms were linked to plant cell walls, extracellular domains, and secretory pathway (**Figure 4B**). We identified several over-represented *PRXs* (15 genes) and EXTs-related proteins (7 encoding genes) suggesting a global change in ROS-homeostasis and EXTs cell wall remodeling in the double *prx62-1 prx69-1* mutant at low-temperature (**Figure 4A-B**). Some of these genes (e.g. *EXT6* and *PRP1*) showed a gene dose-dependent expression at transcript level linked to the RH growth phenotype at 10°C (**Figure 4C**). This indicated that low-temperature induces global gene expression changes linked to the cell wall remodeling and ROS-homeostasis that positively enhance RH growth. The analysis highlights that the absence of PRX62 and PRX69 proteins triggers major changes in the transcriptional program of other PRXs and EXTs genes at low-temperature. This implies the existence of a feedback regulatory loop from the apoplast-cell wall compartments that triggers major changes at the transcriptional level of cell wall proteins and apoplastic PRXs.

**Figure 4.**
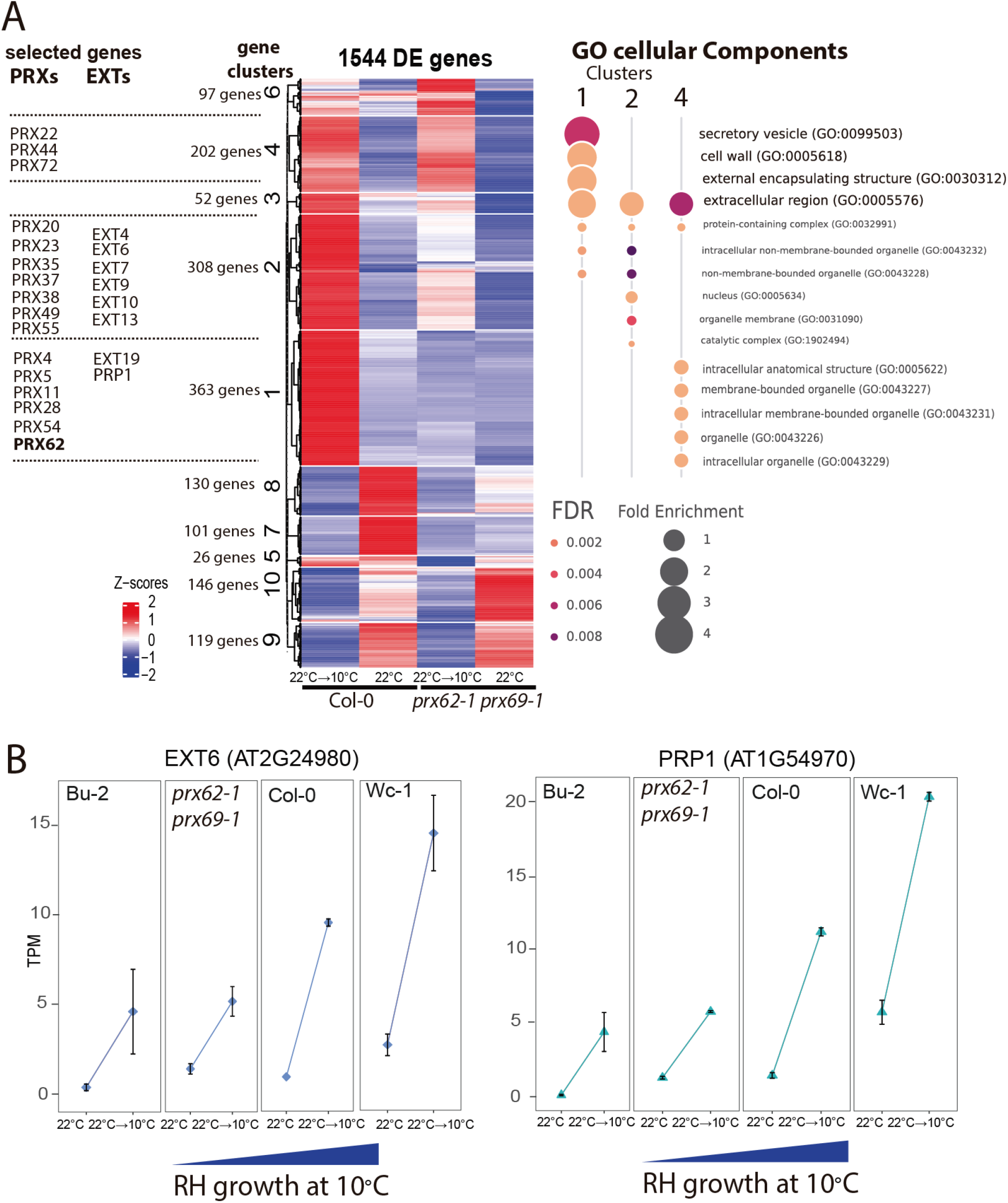
Global transcriptomic changes induced by low-temperature and misregulated in the *prx62 prx69* mutant are associated with PRXs and cell associated-EXTs. (A) Heat-map showing the hierarchical gene clustering for 1,544 *A. thaliana* genes differentially expressed (DE) between room temperature growth (22°C) and low-temperature (10°C) growth in wild type Col-0 and in double mutant *prx62-1 prx69-1* roots. Gene Ontology analysis results depicting the top 7 most significantly enriched GO terms are shown as bubble plots on the right for the clusters of interest. DE genes in clusters 1, 2 and 4 were contrasted against all the expressed genes for GO analysis. The size of the points reflects the amount of gene numbers enriched in the GO term. The color of the points means the p value. Relevant gene examples of specific clusters (1, 2 and 4) are listed on the left. (B) Expression of *EXT6* and *PRP1* is gradually upregulated at low temperature (10°C) in 4 genotypes from very short RHs (Bu-2) to very long RHs (Wc-1) (RNA-seq data). TPM = Transcripts Per Kilobase Million.

### PRX62 and PRX69 affect ROS-homeostasis in RH cells under low-temperature

To get a deeper insight into PRX62 and PRX69 functions in growing RHs at moderate low-temperature, we explored the effect of these PRXs on Reactive Oxygen Species (ROS)-homeostasis. Overall PRX functions are linked to ROS, which are one of the key factors regulating polar growth in RHs (Mangano et al. 2017; Mangano et al. 2018; Marzol et al. 2018). Then, we measured total cytoplasmic ROS (_cyt_ROS) using the cell-permeable fluorogenic probe 2′,7′-dichlorodihydrofluorescein diacetate (H_2_DCF-DA) and apoplastic ROS (_apo_ROS) levels with cell-impermeable Amplex™ UltraRed Reagent in RH tips at 22°C and 10°C (**Figure 5A-B**). The double mutant *prx62-1 prx69-1* showed higher levels of _cyt_ROS at 10°C in actively growing RH tips compared to Col-0 (**Figure 5A**) while this enhance in ROS level is less evident at 22°C between the double mutant *prx62-1 prx69-1* and Col-0. In agreement, in the plants overexpressing PRX62 or PRX69, _cyt_ROS were reduced at both 22°C and 10°C. On the other hand, the _apo_ROS in the RH tip were enhanced in Col-0 at 10°C compared to the levels at 22°C while they were lower in the double mutant *prx62-1 prx69-1* at both temperatures. In the lines overexpressing PRX62 or PRX69, _cyt_ROS were enhanced at both 22°C and 10°C (**Figure 5B**). The increased level of _apo_ROS in Col-0 under low-temperature is in agreement with a two-fold increase in the transcript levels for *NOXC* (*RBOHC*), a key enzyme-encoding gene for ROS production (**Figure S8**). Collectively, these results suggest that ROS-homeostasis is drastically modified in an antagonistic manner by the absence or overexpression of these two PRXs when RH grow at 10°C, affecting RH cell elongation.

**Figure 5.**
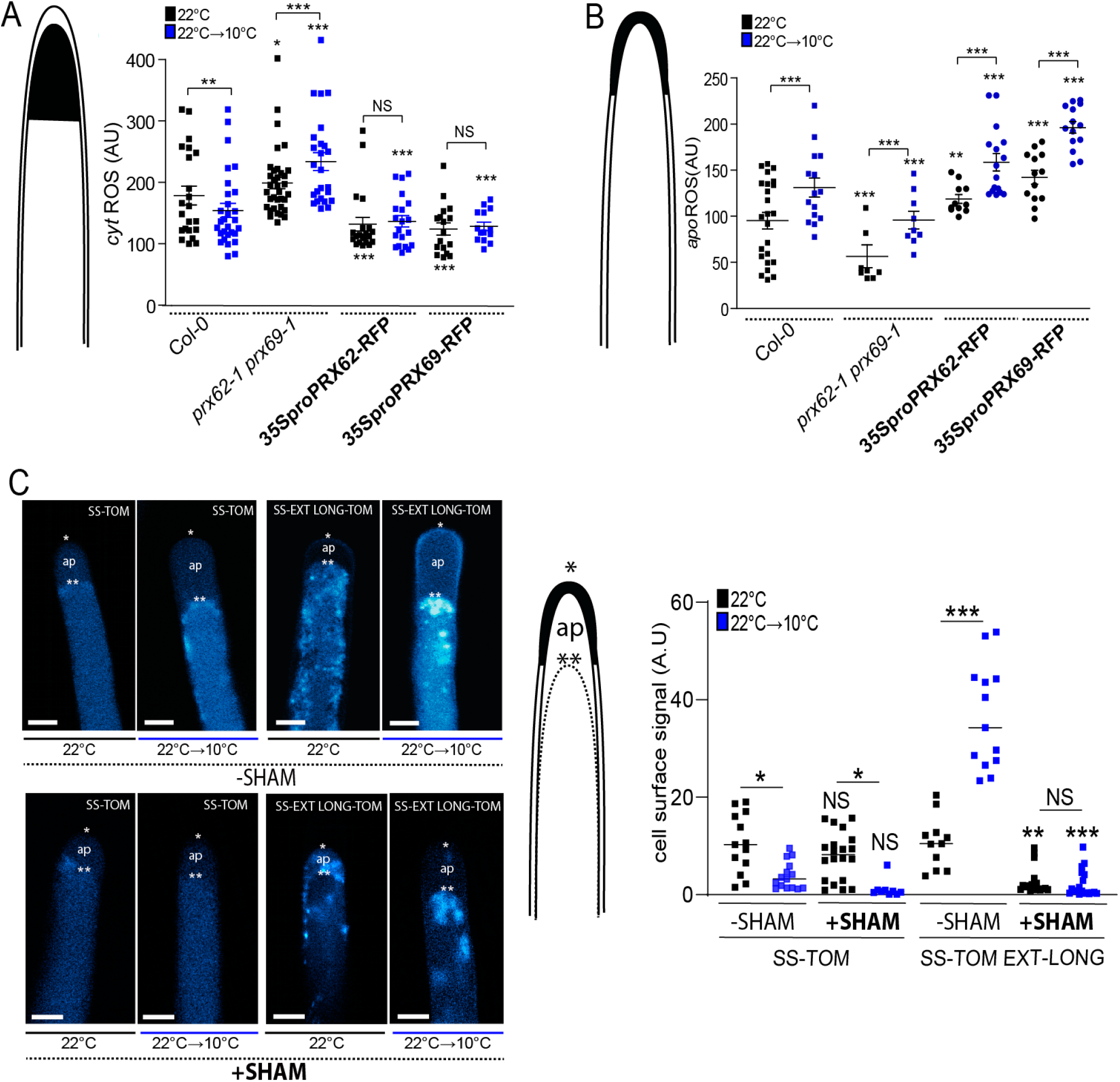
ROS-homeostasis and EXT-stabilization in RH apical cell wall depend on PRX62 and PRX69. (A) Cytoplasmic ROS (_cyt_ROS) levels were measured using 2′,7′-dichlorodihydrofluorescein diacetate (H_2_DCF-DA) in apical areas of RHs in wild-type (Columbia Col-0), in the double mutant *prx62-1 prx69-1* and in the *35S_pro_PRX62*/Col-0 and *35S_pro_PRX69*/Col-0 lines grown at 22°C or 10°C. Measurements are the mean of three replicates ± SD. P-value of one-way ANOVA, (*) P<0.1, (**) P<0.01, (***) P<0.001. NS= non-significant differences. (B) Apoplastic ROS (_apo_ROS) levels were measured with Amplex™ UltraRed in apical areas of RHs in wild-type (Columbia Col-0), in the double mutant *prx62-1 prx69-1* and in the *35S_pro_PRX62*/Col-0 and *35S_pro_PRX69*/Col-0 lines grown at 22°C or 10°C. Measurements are the mean of three replicates ± SD. P-value of one-way ANOVA, (**) P<0.01, (***) P<0.001. (C) Signal of SS-TOM and SS-EXT LONG-TOM in the apical zone of RHs grown at 22°C or 10°C with or without SHAM treatment in Col-0. Cells were plasmolyzed with mannitol 8%. In the images: (*) indicates cell surface including the plant cell walls, (**) indicates the retraction of the plasma membrane, (ap) apoplastic space delimitated between the plant cell wall and the retracted plasma membrane. (A.U.) = Arbitrary Units. Fluorescence AU were reported as the mean of three replicates ± SD. P-value of one-way ANOVA, (*) P<0.05, (**) P<0.01, (***) P<0.001. NS= non-significant differences.

### Low-temperature enhances EXTENSIN cell wall insolubility in RH cells

EXT-crosslinking can provide architectural stabilization for normal wall reinforcement during cell elongation (Srivastava, 2002; Cannon *et al*., 2008; Bashline *et al*., 2014; Bidhendi and Geitmann, 2016; Yaqoob *et al*., 2020). Since changes in ROS-homeostasis could lead to abnormal cell wall secretion and structure, we wondered whether PRX62 and PRX69 might participate in the cell wall glycoprotein EXTs crosslinking during RH growth at low-temperature. Then, we tested if low-temperature could induce a change in the targeting of EXTs secreted and insolubilized in the wall by the activity of these two PRXs. To this end, we used an EXT-reporter carrying a tdTomato tag (SS-TOM-Long-EXT) that is resistant to acidic pH, a condition usually found in the cell wall-apoplast compartments, and a secreted tdTomato tag (SS-TOM) was used as a control (Marzol et al. 2021). The signal coming from the cell surface in the apical zones of RHs cells under plasmolysis conditions were determined for SS-TOM-Long-EXT and SS-TOM constructs at 22°C/10°C temperatures and SHAM-treated/non-treated roots (**Figure 5C**). Plasmolysis allowed us to retract the plasma membrane and define the EXT-signal coming specifically from the cell walls. Interestingly, of cell wall stabilization/insolubility of SS-TOM-Long-EXT in the RH tip was drastically augmented at 10°C compared to 22°C. Furthermore, the signal increment at 10°C was completely abolished when roots were treated with the peroxidase inhibitor SHAM (**Figure 5C**). Thus, the SS-TOM-Long-EXT reporter tested in the apical zone of the RHs is modified by low-temperature and by the peroxidase activity, at least partially possibly exerted by PRX62 and PRX69 in the apoplast. This result suggests that changes in ROS-homeostasis produced by altered levels of these PRXs in the apoplast might affect the secretion, targeting and, possibly the crosslinking of cell wall components including EXTs, affecting RH cell elongation (**Figure 6**).

**Figure 6.**
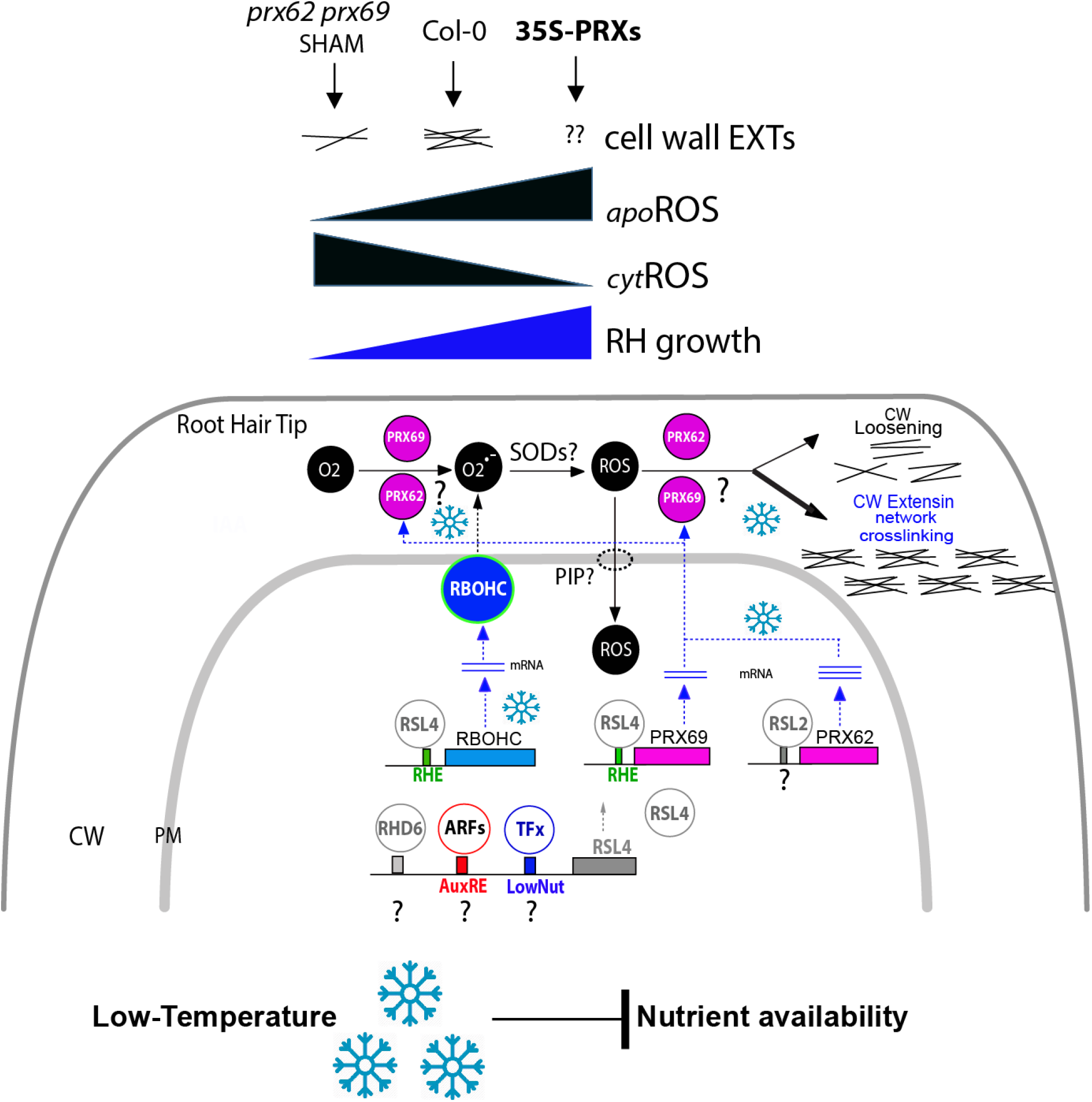
Proposed Model of PRX62 and PRX69 functions in ROS-homeostasis and EXT-cell wall association linked to RH growth at low-temperature. This model is based on the results presented in this study and in previous works (Mangano et al. 2017; Marzol et al. 2021; Moison et al. 2021). ROS-homeostasis (_apo_ROS + _cyt_ROS) in the RH tip. Higher apoROS/low cytROS than Col-0 present in 35S_pro_PRXs promotes RH growth while lower apoROS/high cytROS *in prx62-1 prx69-1* represses RH growth at low-temperature. Changes in _apo_ROS triggered by PRX62 and PRX69 might control changes in the EXT-mediated cell wall expansion/crosslinking in the apical RH zone. Part of the _apo_ROS might be translocated to the cytoplasm to contribute to the _cyt_ROS helped by Plasma membrane Intrinsic Proteins (PIPs). Finally, at the transcriptional level, RSL4 directly regulates *PRX69* expression and indirectly *PRX62* at low-temperature. How RSL4 is regulated under low-temperature remains to be established. Auxin, RHD6, and/or an unknow TF are the most probable regulators of RSL4 expression at low temperature in RHs. AuxRE=Auxin response element. Low-Nut= low nutrient putative *cis*-element; RHE= Root Hair E-box. TFx= unknown TF.

### RSL4 transcription factor binds to the *PRX69* promoter E -boxes

It was previously shown that RSL4 controls RH growth at low-nutrient conditions (Mangano et al. 2017; Mangano et al. 2018) and at low-temperature (Moison et al. 2021). Moreover, it was also shown that RSL4 directly controls the expression of *PRX01*, *PRX44* and *PRX73*, three PRX-encoding genes important for RH growth at low-nutrient condition at room temperature (Mangano et al. 2017; Marzol et al. 2021). Thus, we wondered whether RSL4 was able to regulate directly the expression of *PRX62* and/or *PRX69*. To this end, we first measured *PRX62* and *PRX69* transcript levels in *rsl4* and *rsl2 rsl4* mutants, which is impaired in RH growth enhancement, and in the RSL4 overexpressing line (*35S_pro_RSL4*) that develops extra-long RHs regardless the media and temperature conditions (Moison et al. 2021). PRX62 expression might be positively regulated (indirectly or directly) by RSL2 but not by RSL4 based on the expression profiles in the double mutant *rsl2 rsl4* versus *rsl4* and 35S_pro_RSL4. PRX62 expression enhancement at low temperature is only repressed when *rsl2* mutation is present (**Figure S9A**). On the other hand, an increase of 4.3-folds of *PRX69* transcripts were detected when compared 35S_pro_RSL4/*rsl4* (**Figure S9A**). To test if any of these genes was directly regulated by RSL4, we tested by ChIP-qPCR the binding of RSL4-GFP in the predicted sites (E-boxes) of *PRX62* and *PRX69* promoters using the positive control of *EXPANSIN7* and *LRX1*, two previously reported direct targets of RSL4 (Hwang et al. 2017) and *PP2A* as a negative control (**Figure S9B**). We detected a mild binding of RSL4 protein to one of the predicted E-box sequences in the promoter region *PRX69*. Altogether, our results indicate that RSL4 positively controls the expression of *PRX69* in a direct manner while the regulation of *PRX62* might be related to RSL2. Further research will be required to determine if auxin-ARFs or yet unknown TFs independent of auxin pathway (e.g. RHD6) regulates the expression of *RSL4* under this low-temperature condition. Overall, this work uncovers the key roles of two previously poorly described PRXs, PRX62 and PRX69, in the regulation of low temperature ROS homeostasis and EXT insolubilization in the cell walls that determines an enhanced RH growth. It is hypothesized that these two PRXs might modulate the cell wall EXT-mediated assembly during this fast cell elongation process (**Figure 6**).

## CONCLUSIONS

Despite the putative high overall genetic redundant functions of apoplastic Class-III PRXs, in the last years several individual PRXs were characterized to be involved in the oxidative polymerization of monolignols in the apoplast of the lignifying cells in xylem (e.g. PRX17; Cossio et al 2017), in the root endodermis (e.g. PRX64; Lee et al. 2013) or in petal detachment (Lee et al 2018). Moreover, PRXs are also able to polymerize other components of the plant cell wall, including suberin, pectins and EXTs (Schnabelrauch *et al*., 1996; Bernards et al., 1999; Jackson et al., 2001; Francoz et al. 2019). While several candidates of PRXs have been described in divers plants to be associated specifically to EXTs crosslinking (EXT-PRXs) by *in vitro* studies (LEP1, GvEP1 and FBP1) or inmunolocalization evidences linked them to transient activity measurements (PRX08 and PRX34) (Schnabelrauch et al., 1996; Wojtaszek et al., 1997; Jackson et al., 2001; Price et al., 2003; Pereira et al. 2011; Dong et al., 2015; Jacobowitz et al. 2019), their role *in vivo* remains largely unexplored. Previously it was demonstrated that three PRXs (PRX01, PRX44 and PRX73) directly contribute to ROS-homeostasis and RH growth at room temperature (22°C) under low-nutrient condition (Mangano et al. 2017; Marzol et al. 2021). By using a GWAS-RNAseq approach, we identified here two previously poorly characterized apoplastic peroxidases, PRX62 and PRX69 (Jemmat et al. 2020), as positive regulators of RH growth at low-temperature (10°C). One of the key results of this work is that PRX62 was found using GWAS while PRX69 was identify on the transcriptomic profile. This point out to a different evolutionary history for both proteins. PRX62 has evolved to give a dose response according to the allele encoded in the genome, while PRX69 have a constitutive response at low temperature. These features of PRX62 and PRX69 can be useful in crop improvement, to select varieties with differential responses; better adapted to the environment they are exposed. The evidences shown here indicate that PRX62 and PRX69 are involved in the ROS-homeostasis linked to the association of EXTs to the cell wall during RH cell elongation at low-temperature (**Figure 6**). We speculate that cell wall insolubilization/association of EXTs triggered by low-temperature might not only involve Tyr-covalent crosslinks mediated by these two PRXs identified here but also by EXT hydrophobic associations non-dependent on Tyr as suggested before for Leucine-Rich Extensins 1 (LRX1; Ringli 2010). Further analyses might shed light on these complex processes.

Previously, our group as well as others have documented that changes in any of the several posttranslational modifications in EXTs and related-EXTs like LRXs (e.g. proline hydroxylation, *O*-glycosylation, and Tyr-crosslinking), all affected RH growth (Baumberge et al. 2001, 2003; Velasquez et al. 2011; Velasquez et al. 2015; Marzol 2018, 2021) as well as pollen tube growth (Fabrice et al. 2017; Sede et al. 2017; Wang et al. 2017). In addition, auxin-dependent ROS-homeostasis controlled by three apoplastic PRXs (e.g. PRX01, PRX44, PRX73) and plasmamembrane RBOHC protein (also known as RHD2, for RH Defective 2) was shown to be determinant for a proper RH growth under low nutrient condition (Mangano et al 2017; Marzol et al. 2021) or under low temperature (Martinez-Pacheco et al. 2021). Collectively, these evidences highlight the predominant role of ROS-homeostasis partially regulated by specific PRXs as a key component in polar RH elongation. The molecular mechanism by which low-temperature-associated nutrient availability in the media (Moison et al. 2021) triggers the expression of these two specific PRXs remains unclear, although RSL4 could play a central role in the regulation of this mechanism. Previously, we have shown that the lncRNA *APOLO* binds to the locus of RHD6 and controls *RHD6* transcriptional activity leading to cold-enhanced RH elongation through the consequent activation of *RSL4* (Moison et al. 2021) and of several cell wall EXTENSIN (EXT) encoding genes (Martinez-Pacheco et al. 2021). Unexpectedly, our previous results indicate that the low-temperatures (10°C) are able to trigger an exacerbated RH growth compared with cell expansion at room temperature (Moison et al. 2021; Martinez-Pacheco et al. 2021). Moreover, further research will be needed to uncover the nutritional signal perceived at the RH cell surface to trigger PRX62 and PRX69 low temperature mediated growth response. The expression levels of PRX62 and PRX69 orthologs in other Brassicaceae could be used as biomarkers for crop improvement in the selection of genotypes with longer RHs at moderate low-temperatures in order to boost nutrients uptake in deficient soils.

## EXPERIMENTAL PROCEDURES

### Plant genotyping and growth conditions

*Arabidopsis thaliana* Columbia-0 (Col-0) was used as the wild type (Wt) genotype in all experiments unless stated otherwise. Seedlings were surface sterilized and stratified in darkness at 4°C for 3 days before been germinated on ½ strength MS agar plates supplemented with MES (Duchefa, Netherlands), in a plant growth chamber in continuous light (120 µmol.sec^−1^.m^−2^). Plants were transferred to soil for growth under the same conditions as previously described at 22°C. Mutants and transgenic lines developed and used in this study are listed in **Table S3**. For identification of T-DNA knockout lines, genomic DNA was extracted from rosette leaves. Confirmation by PCR of a single and multiple T-DNA insertions in the genes were performed using an insertion-specific LBb1 or LBb1.3 (for SAIL or SALK lines, respectively) or 8474 (for GABI line) primer in addition to one gene-specific primer. In this way, we isolated homozygous for all the genes. *Arabidopsis* T-DNA insertions lines (*prx62-1* [GK_287E07], *prx62-2* [SALK_151762], *prx69-1* [SAIL_691_G12], *prx69-2* [SALK_137991]) were obtained from the European Arabidopsis Stock Centre (http://arabidopsis.info/). Using standard procedures homozygous mutant plants were identified by PCR genotyping with the gene-specific primers listed in **Table S4**. T-DNA insertion sites were confirmed by sequencing using the same primers. Plants were routinely grown in Jiffy peat pellets (continuous light, 120 μmol photons/m/s, 22°C, 67% relative humidity). For *in vitro* experiments, seeds were surface-sterilized and sown in Petri dishes on agar-solidified half-MS medium without sucrose, and grown in a culture room with continuous light (120 μmol photons/m/s, 22°C).

### Root hair phenotype

Seeds were surface sterilized and stratified in darkness for 3 days at 4°C. Then grew on ½ strength MS agar plates supplemented with MES (Duchefa, Netherlands), in a plant growth chamber at 22°C in continuous light (120 µmol.sec^−1^.m^−2^) for 5 days at 22°C + 3 days at 10°C or for 8 days at 22°C as *control*. For quantitative analysis of root hair cell length phenotypes, 10 fully elongated RHs from the elongation root zone were measured from 15-20 roots. Measurements were made after 8 days. Images were captured with an Olympus SZX7 Zoom Stereo Microscope (Olympus, Japan) equipped with a Q-Colors digital camera and QCapture Pro 7 software (Olympus, Japan) and digitally processed with ImageJ software. RH length values were reported as the mean of three replicates ± SD using the GraphPad Prism 8.0.1 (USA) statistical analysis software.

### GWAS analysis and haplotype analysis

To perform Genome Wide Association Analysis (GWAS), 106 *Arabidopsis thaliana* natural accessions were phenotyped for RH length in a shift-temperature experiment as described above (**Table S1**). The population was previously genotyped using 214,051 Single Nucleotide Polymorphisms (SNPs) and this information is publicly available (Horton et al., 2012). These set of phenotypes and genotypes were used to performed GWAS on the GWAPP web application from the GWA-Portal (Seren et al, 2012, https://gwas.gmi.oeaw.ac.at/#/home, Experiment code: 3b316208-0b5d-11e7-b6b1-005056990049) applying the accelerated mixed model, AMM (Kang et al., 2008; Zhang et al., 2010, Kang et al. 2010). A total of 139,425 SNPs with minor allele frequency (MAF) ≥10% were retained for further analysis. P-values of association were log-transformed to LOD values (–log_10_ (p-value)) and corrected for multiple comparisons using FDR procedure (Benjamini and Hochberg, 1995). The threshold for significant associations was set to p-value ≤ 1/N (where N is the number of SNPs= 139,425) as described previously (Wen et al. 2014). Manhattan plots were obtained using the qqman package (Turner, 2017) in R (2013), filtering out the SNPs with p-value > 0.4, to minimize overrepresentation of non-significant SNPs. Linkage disequilibrium, i.e. the degree to which an allele of one SNP co-occurs with an allele of another SNP within a population, was calculated as square coefficient of correlation (r^2^) and visualized using the LDheatmap package (Shin et al., 20006) in R. Three additional SNPs in the *PRX62* genomic region (m190904, m190907, m190909) in high linkage disequilibrium (r^2^ > 0.7, p << 0.001) with the lead SNP m190905 were used in the haplotype analysis. Mean trait values for each non-unique haplotype were analyzed using ANOVA followed by Tukey test implemented in Infostat (Di Rienzo et al. 2011).

### Peroxidase activity

Soluble proteins were extracted from roots grown on vertical plates for 10 days at 22°C or 10°C by grinding in 20mM HEPES, pH 7.0, containing 1 mM EGTA, 10mM ascorbic acid, and PVP PolyclarAT (100mg/g fresh material)(Sigma, Buchs, Switzerland). The extract was centrifuged twice for 10 min at 10,000 g. Each extract was assayed for protein levels with the Bio-Rad protein assay (Bio-Rad, USA). Enzyme activity (expressed in nkatal/mg protein) was determined at 25°C by following the oxidation of 8 mM guaiacol (Fluka^TM^, Honeywell International,USA) at 470 nm in the presence of 2 mM H_2_O_2_ (Carlo Erba, Italy) in a phosphate buffer (200 mM, pH6.0). Values are the mean of three replicates ± SD. P-*value* of one-way ANOVA, (**) P<0.01.

### Gene transcript analysis by Reverse Transcription followed by quantitative PCR (RT-qPCR)

Total RNA was prepared from 10 days old *in vitro*-grown plantlets using the TRI^TM^ Reagent Solution (Sigma-Aldrich). After quantification by spectrophotometry and verification by electrophoresis, RNA was treated with the RQ1 RNase-free DNase I (Promega). One microgram of total RNA was reverse transcribed using an oligo(dT)_15_ and the MMLV-RT (Promega) according to the manufacturer’s instructions. cDNA was diluted 20-fold before PCR. RT-qPCR were performed on a QuantStudio 6 Flex Real-Time PCR System (Thermo Fisher) using 5 μL Power SYBR Green PCR Mix (Applied Biosystems), 2 μL of cDNA, and 0.3 μM of each primer in a total volume of 10 μL per reaction. Primers used are listed in **Table S4**. ACT2 (AT3G18780) and UBQ1 (AT3G52590) genes were used as references for normalization of gene expression levels. The cycling conditions were 95°C for 10 min., 40 cycles of 95°C for 15 sec., 60°C for 1 min. and finally a melting curve from 60°C to 95°C (0.05°/sec). Under these conditions primers efficiency was found to be between 97.0 and 99.7%. No amplification occurred in the no-template controls. Data were analyzed using the Standard curve method (Pfaffl, 2001) and Qiagen REST© 2009 software (Pfaffl et al., 2002). Three independent experiments (and two technical replicates per experiment) were performed.

### PRXs-tagged reporter lines

For the PRX62proGFP and PRX69proGFP reporter lines, a 1.5 kb genomic region upstream of the ATG start codon of each PRX62 (AT5G39580) and PRX69 (AT5G64100) genes was selected using ThaleMine (https://bar.utoronto.ca/thalemine/begin.do) synthetized and cloned into the pUC57 vector by GenScript Biotech(USA). Through Gateway cloning Technology (Invitrogen) the 1.5 kb upstream regions were recombined first in pDONR™207 vector and subcloned into pMDC111 destination vector (Invitrogen; (Karimi et al., 2002)) for PRX69 promoter region and into pGWB4 vector (Invitrogen; (Nakagawa et al 2007)) for PRX62 promoter region. These constructs were checked by restriction analysis. Both vectors contain a cassette with a C-terminal GFP tag. For the PRXs-TagRFP lines, the PRX62 and PRX69 coding sequence was amplified by PCR from *A. thaliana* 10-day old plantlets cDNAs using specific primers (**Table S4**). The PCR product was digested with *Hind*III and *Bam*HI (PRX62) or with *Eco*RI and *Sma*I (PRX69), and cloned into Gateway® TagRFP-AS-N entry clone (Evrogen). The PRX62-TagRFP fusion was subcloned (Gateway Technology, Invitrogen) into the pB7WG2 destination vector (Karimi et al., 2002) containing a 35S promoter. This construct was checked by restriction analysis and sequencing. The same procedure was use to generate 35S-PRX69-TagRFP construct. All the constructs were used to transform *A. thaliana* plants and obtain homozygous stable fluorescent lines.

### Confocal Microscopy

Confocal laser scanning microscopy for the lines *PRX62pro-GFP* and *PRX69proGFP,* was performed using Zeiss LSM5 Pascal (Zeiss, Germany) (Excitation: 488 nm argon laser; Emission: 490-525 nm, 10X objective N/A 0.30 or 40X water-immersion objective, N/A 1.2, according to experiment purpose). Z stacks were done with an optical slice of 1µm, and fluorescence intensity was measured at the RH tip. For the lines *SS-TOMATO* and *SS-TOMATO-EXT LONG*, roots were plasmolyzed with a mannitol 8% solution and the scanning was performed using Zeiss LSM5 Pascal (Zeiss, Germany)( Excitation: 543 nm argon laser; Emission: 560-600 nm, 63X water-immersion objective N/A 1.2) Three replicates for each of ten roots and between 10 to 15 hairs per root were observed. GFP signal and tdTOMATO cell wall signal at RH tip were quantified using the ImageJ software. Fluorescence AU were reported as the mean of three replicates ± SD using the GraphPad Prism 8.0.1 (USA) statistical analysis software.

### Apoplastic and Cytoplasmic ROS measurement in RH Tip

To measure ROS levels in root hairs cells, 8 days-old Arabidopsis seedlings grown at 22°C (control) and 10°C in continuous light were used. For cytoplasmic ROS, the seedlings were incubated in darkness for 10 min with 50 μM H2DCF-DA (Thermo Fisher) at room temperature then washed with liquid 0.5X MS media (Duchefa, Netherlands) and observed with Zeiss Imager A2 Epifluorescence Microscope(Zeiss, Germany) (40X objective, N/A 1.2, exposure time 25 ms). Images were analyzed using ImageJ software. To measure ROS levels, a circular region of interest was chosen in the zone of the root hair tip cytoplasm. Approximately, 20 healthy and alive root hairs of ten plants per treatment were analyzed. To measure apoplastic ROS, the seedlings were incubated with 50 µM Amplex™ UltraRed Reagent (AUR) (Molecular Probes, Invitrogen) for 15 min in darkness and rinsed with liquid 0.5X MS media (Duchefa, Netherlands). Root hairs were imaged with a Zeiss LSM5 Pascal (Zeiss, Germany)) laser scanning confocal microscope (Excitation: 543 nm argon laser; Emission: 560-610 nm, 40X water-immersion objective, N/A 1.2). The intensity of fluorescence was quantified on digital images using ImageJ software. Quantification of the AUR probing fluorescence signal was restricted to apoplastic spaces at the root hair tip. At least 10-15 hairs per plant and ten plants per treatment with three replicates were analyzed. Fluorescence AU were reported as the mean of three replicates ± SD using the GraphPad Prism 8.0.1 (USA) statistical analysis software.

### In silico analysis

The *in silico* analysis of *PRX62* and *PRX69* expression in the roots were performed using ePlant browser of Araport, Tissue Specific Root eFP (http://bar.utoronto.ca/eplant/) (Waese et al. 2017). EXPANSIN7 (EXP7) as a RH specific gene was included for comparison.

### RNA-seq Analyses

This section is adapted from the 3D RNA-seq package output “Results” (Guo et al., 2020; Calixto et al., 2018) as this was the selected tool to analyze differential expression in our datasets. For the RNA-seq datasets we analyzed 2 datasets, one with 16 factor groups (Col.X10, Col.X22, Bu.X10, Bu.X22, Sf.X10, Sf.X22, Te.X10, Te.X22, Wc.X10, Wc.X22, P62.X10, P62.X22, P69.X10, P69.X22, P6269.X10, P6269.X22) each with 2 biological replicates (32 samples in total). Quantification of transcripts using salmon quant (Patro et al., 2017) from Galaxy.org or salmon-1.5.1_linux_x86_64 version in a personal computer. The index of the transcriptome was built using The Arabidopsis Thaliana Reference Transcript Dataset 2 (AtRTD2, Zhang R. et al., 2016) obtained from https://ics.hutton.ac.uk/atRTD/. For the data pre-processing, read counts and transcript per million reads (TPMs) were generated using tximport R package version 1.10.0 and lengthScaledTPM method (Soneson et al., 2016) with inputs of transcript quantifications from tool salmon (Patro et al., 2017). Low expressed transcripts and genes were filtered based on analyzing the data mean-variance trend. The expected decreasing trend between data mean and variance was observed when expressed transcripts were determined as which had ≥ 1 of the 32 samples with count per million reads (CPM) ≥ 1, which provided an optimal filter of low expression. A gene was expressed if any of its transcripts with the above criteria was expressed. The TMM method was used to normalize the gene and transcript read counts to *log*_2_-CPM (Bullard et al., 2010). The principal component analysis (PCA) plot showed the RNA-seq data did not have distinct batch effects. For the DE, DAS and DTU analysis, the voom pipeline of limma R package was used for 3D expression comparison (Ritchie et al., 2015; Law et al., 2014). To compare the expression changes between conditions of experimental design, the contrast groups were initially set as Col.X10-Col.X22, Bu.X10-Bu.X22, Sf.X10-Sf.X22, Te.X10-Te.X22, Wc.X10-Wc.X22, P62.X10-P62.X22, P69.X10-P69.X22, P6269.X10-P6269.X22. For DE genes/transcripts, the *log*_2_ fold change (*L*_2_*FC*) of gene/transcript abundance were calculated based on contrast groups and significance of expression changes were determined using t-test. P-values of multiple testing were adjusted with BH to correct false discovery rate (FDR) (Benjamini and Yekutieli, 2001). A gene/transcript was significantly DE in a contrast group if it had adjusted p-value < 0.01 and *L*_2_*FC* ≥ 1.5. Heatmap: Hierarchical clustering was used to partition the DE genes into 10 clusters with euclidean distance and ward.D clustering algorithm (Saracli et al., 2013). ComplexHeatmap R package version 1.20.0 was used to make the heat-map.

### Alternative Splicing analysis

At the alternative splicing level, DTU transcripts were determined by comparing the *L*_2_*FC* of a transcript to the weighted average of *L*_2_*FCs* (weights were based on their standard deviation) of all remaining transcripts in the same gene. A transcript was determined as significant DTU if it had adjusted p-value < 0.01 and *Δ*PS ≥ 0.15. For DAS genes, each individual transcript *L*_2_*FC* were compared to gene level *L*_2_*FC*, which was calculated as the weighted average of *L*_2_*FCs* of all transcripts of the gene. Then p-values of individual transcript comparison were summarized to a single gene level p-value with F-test. A gene was significantly DAS in a contrast group if it had an adjusted p-value < 0.01 and any of its transcript had a *Δ* Percent Spliced (*Δ*PS) ratio ≥ 0.15.

### Gene Ontology Analysis

Gene ontology (GO) terms assignment for the DE genes datasets were obtained using the PantherDB tool (http://go.pantherdb.org/index.jsp). An enrichment test was performed for the following categories: BP (biological process), MF (molecular function), and CC (cellular component). *p*-values were obtained using the Fisher exact test and corrected for multiple testing using FDR. The enrichment factor (EF) was estimated as the ratio between the proportions of genes associated with a particular GO category present in the dataset under analysis, relative to the number of genes in this category in the whole transcriptome analyzed. We considered the whole transcriptome as those genes that are expressed at least in one of the evaluated conditions. Bubble plots were generated, using a custom script written in Python language (https://github.com/Lucas-Servi/makeGO), for all those categories for which the adjusted p-value (FDR) was lower than 0.01.

### Chromatin immunoprecipitation (CHIP) assays

ChIP assays were performed on seedlings expressing RSL4-GFP under the native RSL4 promoter, using anti-GFP (Abcam ab290) and anti-IgG (Abcam ab6702) antibodies, mainly as described by Ariel et al. (2020). Seedlings were grown in continuous light at 22°C for 10 days, harvested after 24 h treatment at 10°C, ground in liquid nitrogen, and resuspended in 25 mL of nuclear isolation buffer (Bourbousse et al. 2018). Chromatin was cross-linked first with 1.5 mM ethylene glycol bis(succinimidyl succinate) (Thermo Fisher Scientific 21565) for 20 min at room temperature, and then with formaldehyde at a final concentration of 1% for 10 min at room temperature. Cross-linking was stopped by adding 1.7 mL of 2 M glycine and incubating for 10 min at room temperature. Crosslinked chromatin was extracted by cell resuspension, centrifugation, cell membrane lysis, and sucrose gradient as previously described (Ariel et al., 2020). Nuclei were resuspended in Nuclei Lysis Buffer and chromatin was sonicated using a water bath Bioruptor Pico (Diagenode; 30 s on / 30 s off pulses, at high intensity for 10 cycles). Chromatin samples were incubated for 12 h at 4 °C with Protein A Dynabeads (Invitrogen) precoated with the corresponding antibodies. Immunoprecipitated DNA was recovered using Phenol:Chloroform:Isoamilic Acid (25:24:1; Sigma) and analyzed by qPCR. Untreated sonicated chromatin was processed in parallel and considered the Input sample.

### SHAM Treatment

Seeds were germinated on agar plates at 22°C in a growth chamber in continuous light. After 4 days plants were transferred to agar plates with or without 65 µM of SHAM (Salicylhydroxamic acid; Sigma Aldrich, USA), then grown 3 days at 22°C followed by 3 days at 10°C or 6 days at 22°C (control). Root hair phenotype was measured and confocal microscopy analysis was performed.

### SS-TOM and SS-TOM-Long-EXT constructs

The binary vector pART27, encoding tdTomato secreted with the secretory signal sequence from tomato polygalacturonase and expressed by the constitutive CaMV 35S promoter (pART-SS-TOM), was a kind gift of Dr. Jocelyn Rose, Cornell University. The entire reporter protein construct was excised from pART-SS-TOM by digesting with *Not*I. The resulting fragments were gel-purified with the QIAquick Gel Extraction Kit and ligated using T4 DNA Ligase (New England Biolabs) into dephosphorylated pBlueScript KS+, also digested with *Not*I and gel-purified, to make pBS-SS-TOM. The plasmid was confirmed by sequencing with primers 35S-FP (5’-CCTTCGCAAGACCCTTCCTC-3’) and OCS-RP (5’-CGTGCACAACAGAATTGAAAGC-3’). The sequence of the EXT domain from *SlPEX1* (NCBI accession AF159296) was synthesized and cloned by GenScript into pUC57 (pUC57-EXT). The plasmid pBS-SS-TOM-Long-EXT was obtained by digesting pUC57-EXT and pBS-SS-TOM with *Nde*I and *SgrA*I, followed by gel purification of the 2243 bp band from pUC57-EXT and the 5545 bp band from pBS-SS-TOM, and ligation of the two gel-purified fragments. The pBS-SS-TOM-Long-EXT plasmid was confirmed by sequencing with 35S-FP, OCS-RP, and tdt-seq-FP (5’-CCCGTTCAATTGCCTGGT-3’). Both pBS plasmids were also confirmed by digestion. The binary vector pART-SS-TOM-Long-EXT was made by gel purifying the *Not*I insert fragment from the pBS-SS-TOM-Long EXT plasmid and ligating it with pART-SS-TOM backbone that had been digested with NotI, gel purified, and dephosphorylated. This plasmid was confirmed by sequencing. The construct SS-TOM and SS-TOM-Long-EXT where transformed into Arabidopsis plants. The secretory sequence (SS) from tomato polygalacturonase is MVIQRNSILLLIIIFASSISTCRSGT (2.8kDa) and the EXT-Long domain sequence is BAAAAAAACTLPSLKNFTFSKNIFESMDETCRPSESKQVKIDGNENCLGGRSEQRTEKECFPVVSKPVDCSKGHCG VSREGQSPKDPPKTVTPPKPSTPTTPKPNPSPPPPKTLPPPPKTSPPPPVHSPPPPPVASPPPPVHSPPPPVASPPPPVHSPPPPPVASPPPPVHSPPPPVASPPPPVHSPPPPVHSPPPPVASPPPPVHSPPPPVHSPPPPVHSPPPPVHSPPPPVHSPPPPVASPPPPVHSPPPPVHSPPPPVHSPPPPVASPPPPVHSPPPPPPVASPPPPVHSPPPPVASPPPPVHSPPPPVASPPPPVHSPPPPVHSPPPPVHSPPPPVASPPPALVFSPPPPVHSPPPPAPVMSPPPPTFEDALPPTLGSL YASPPPPIFQGY* 395-(39.9kDa). The predicted molecular size for SS-TOM protein is 54.2 kDa and for SS-TOM-EXT-Long Mw is 97.4 kDa.

## Acknowledgements

We would like to thank Margaret Fleming and Patricia Bedinger for sharing SS-TOM-Long-EXT and SS-TOM constructs and Jorge Muschietti for his valuable comments on this work. We thank NASC (Ohio State University) for providing T-DNA lines seed lines. J.M.E., C.M.F., E.P., J.B are investigators of the National Research Council (CONICET) from Argentina. This work was supported by grants from ANPCyT (PICT2017-0066, and PICT2019-0015 to J.M.E). In addition, this research was also funded by ANID – Programa Iniciativa Científica Milenio ICN17_022 and Fondo Nacional de Desarrollo Científico y Tecnológico [1200010] to J.M.E.

## Author Contribution

J.M.P. performed most of the experiments and analysed the data. P.R. measured the peroxidase activity and PRXs expression in single *prx* mutants and selected accessions, analysed the subcellular localization of PRXs, cloned the PRXs, generated 35SproPRXs constructs, produced the RNA-seq data. L.K performed GWAS measurements. C.M.F. performed GWAS and haplotype analysis. L.S., R.T, C.M, and E.P. performed the bioinformatics analysis. V.B.G., J.M.P., C.B., E.M., D.R.R.G, Y.R., M.C., analysed part of the data. J.B and M.Y. provided the accessions and analysed the data. L.S, P.T., E.P., C.M. analysed the RNA-seq data. In addition, L.S and E.P. analysed the alternative splicing. L.F. and F.A. performed the CHIP assay. C.D. produced the RNA-seq data and analysed the results. J.M.E. designed research, analysed the data, supervised the project, and wrote the paper. All authors commented on the results and the manuscript. This manuscript has not been published and is not under consideration for publication elsewhere. All the authors have read the manuscript and have approved this submission.

---

**Figure S1.**
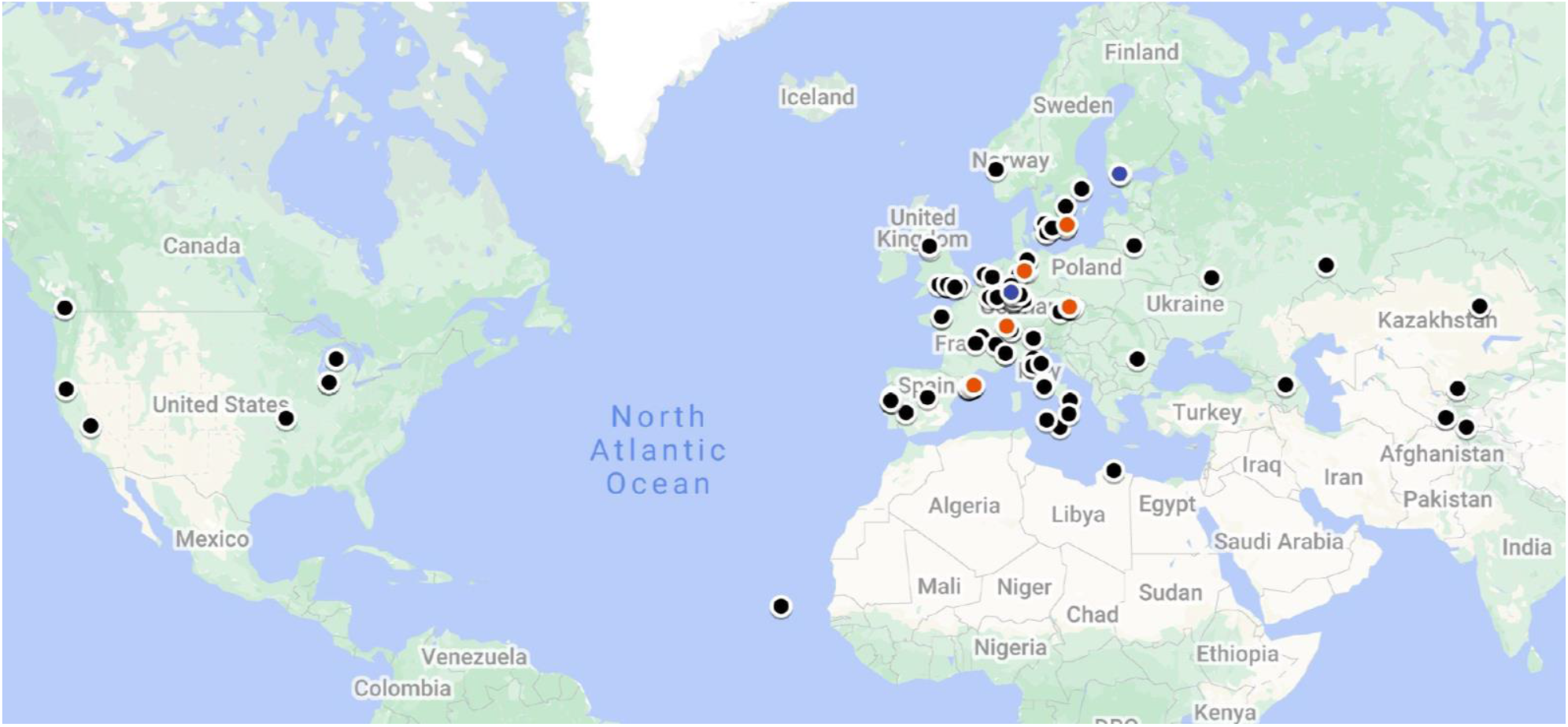
Geographic location of the 106 *Arabidopsis thaliana* accessions used in this study. Each dot represents the sampling original site of individual accession used for this study. In red color, 5 accessions with the longest RHs at 10°C, and in blue those with the shortest RHs at 10°C.

**Figure S2.**
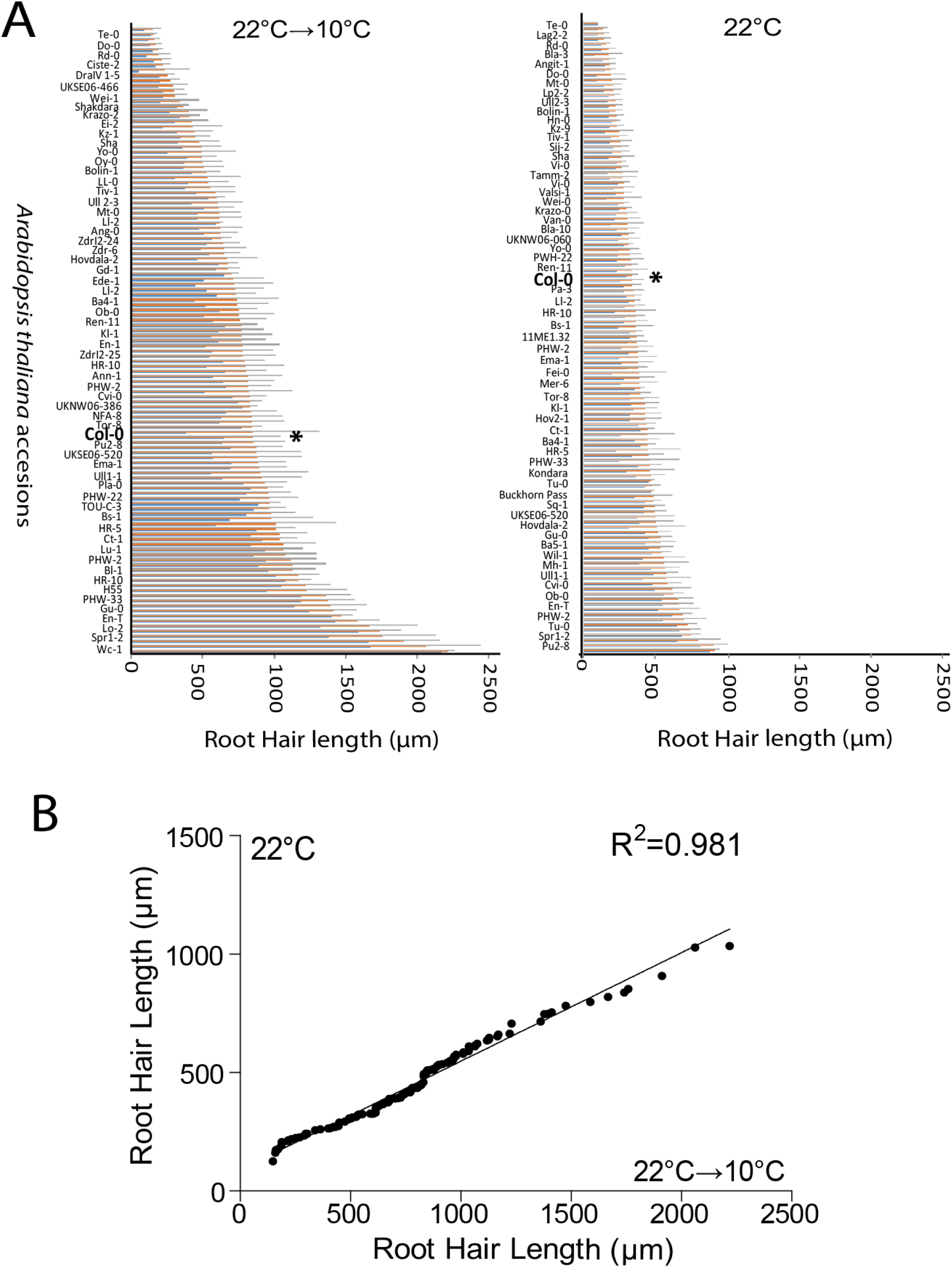
RH cell growth phenotype in *Arabidopsis* accessions at 22°C→10°C versus at 22°C. (A) RH length phenotype in the *Arabidopsis* accessions at 22°C→10°C versus at 22°C. Average cell length on 50-300 fully elongated RHs is indicated (± SD) (N=root 5-30). Col-0 is indicated with an asterisk (*). Only 61 accessions are indicated in the edited graphs to improve readability. Average cell length (in orange), highest (in grey) and lowest (blue) values are shown. (B) Pearson correlation (R^2^=0.981) between RH growth of *Arabidopsis* accessions grown at 22°C versus the same accessions grown at 22°C→10°C.

**Figure S3.**
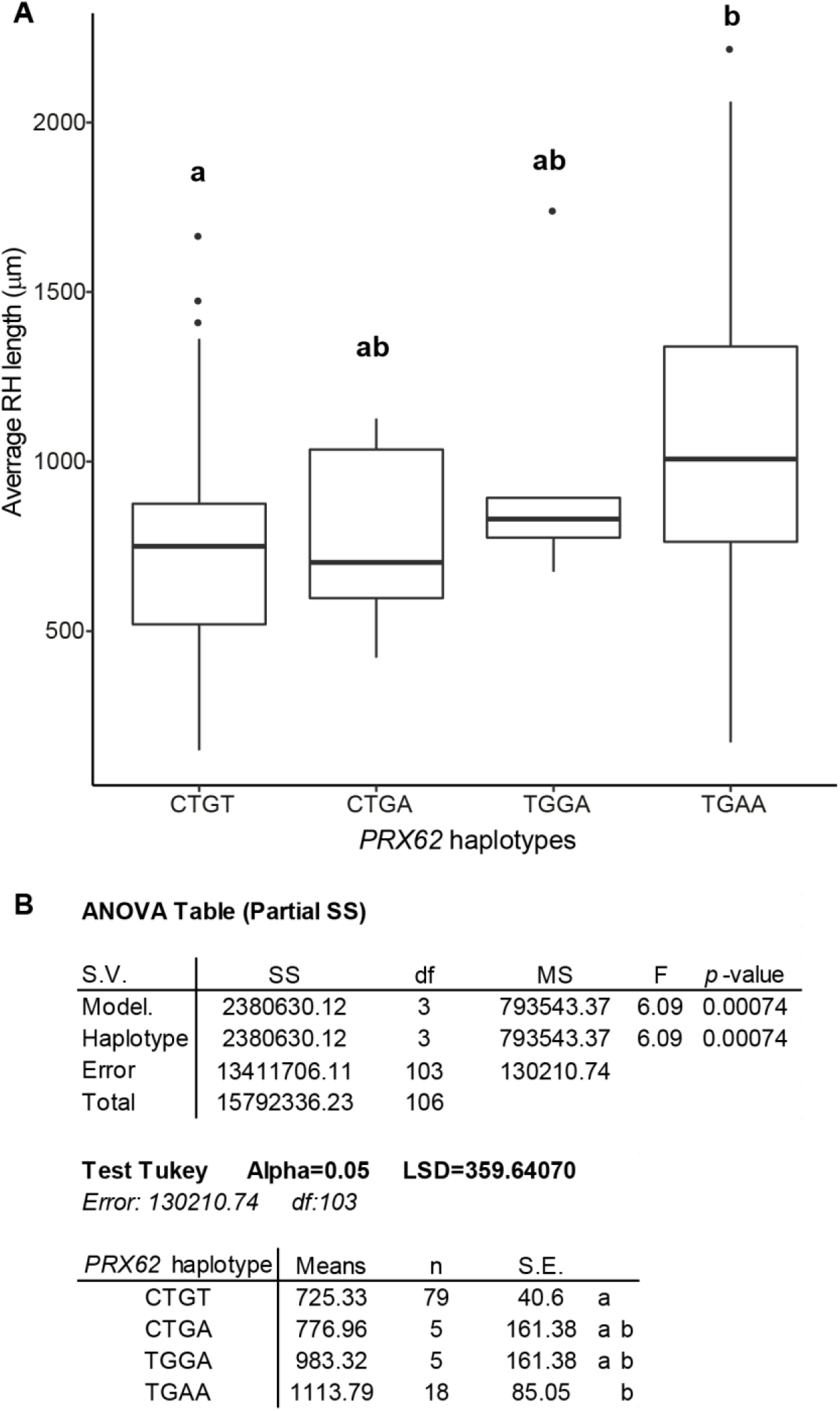
Haplotype analysis on *PRX62* SNPs. (A) Average RH length at 10°C was calculated for each informative haplotype (number of accessions carrying the haplotype ≥ 5) obtained with the four highly associated SNPs identified by GWAS and localized in *PRX62*. Significant differences are indicated by different letters above each haplotype. (B) Model details and contrast for one-way ANOVA. Haplotype contrasts were identified in a post-hoc Tukey HSD test (p ≤ 0.05). Significant differences are indicated by different letters.

**Figure S4.**
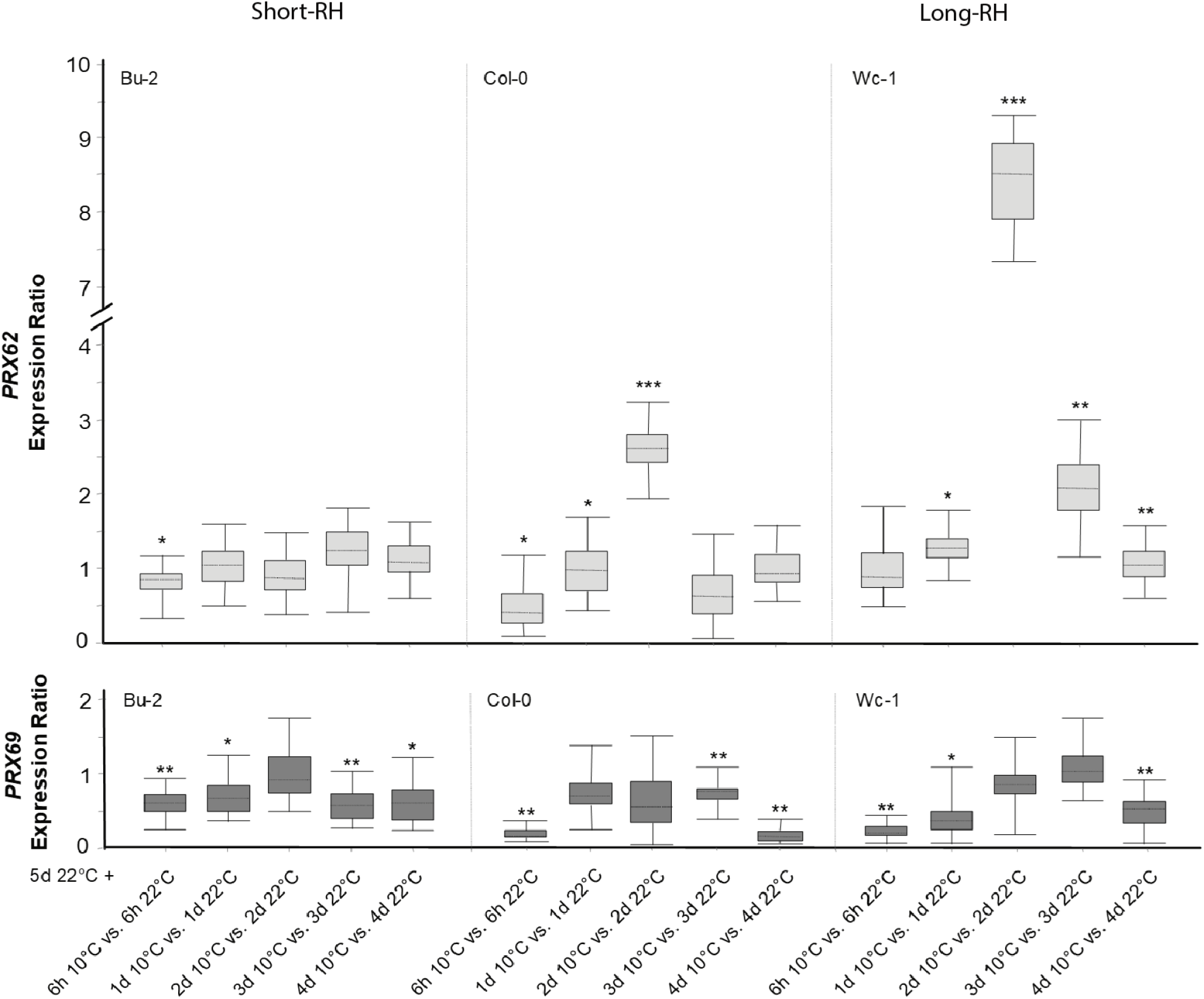
*PRX62,* but not *PRX69,* is differentially expressed at low-temperature (10°C) in contrasting Arabidopsis accessions with contrasting RH phenotypes. Expression measured by qPCR of *PRX62* and *PRX69* in three contrasting Arabidopsis accessions based on the RH phenotype detected at 10°C. Total RNA was extracted from roots of *in vitro* plantlets (grown for 5 days at 22°C plus 6h, 1 day, 2 days, 3 days or 4 days either at 22°C or 10°C). *PRX62* and *PRX69* transcript levels determined by RT-qPCR were normalized to *ACT2* and *UBQ1* as internal controls. Boxes represent the interquartile range. The dotted line symbolizes the median gene expression. Whiskers correspond to the minimum and maximum observations (n = 6). Asterisks indicate statistically significant differences between cold-treated and non-cold-treated groups (***) P<0.001, (**)P<0.01, (*)P<0.05.

**Figure S5.**
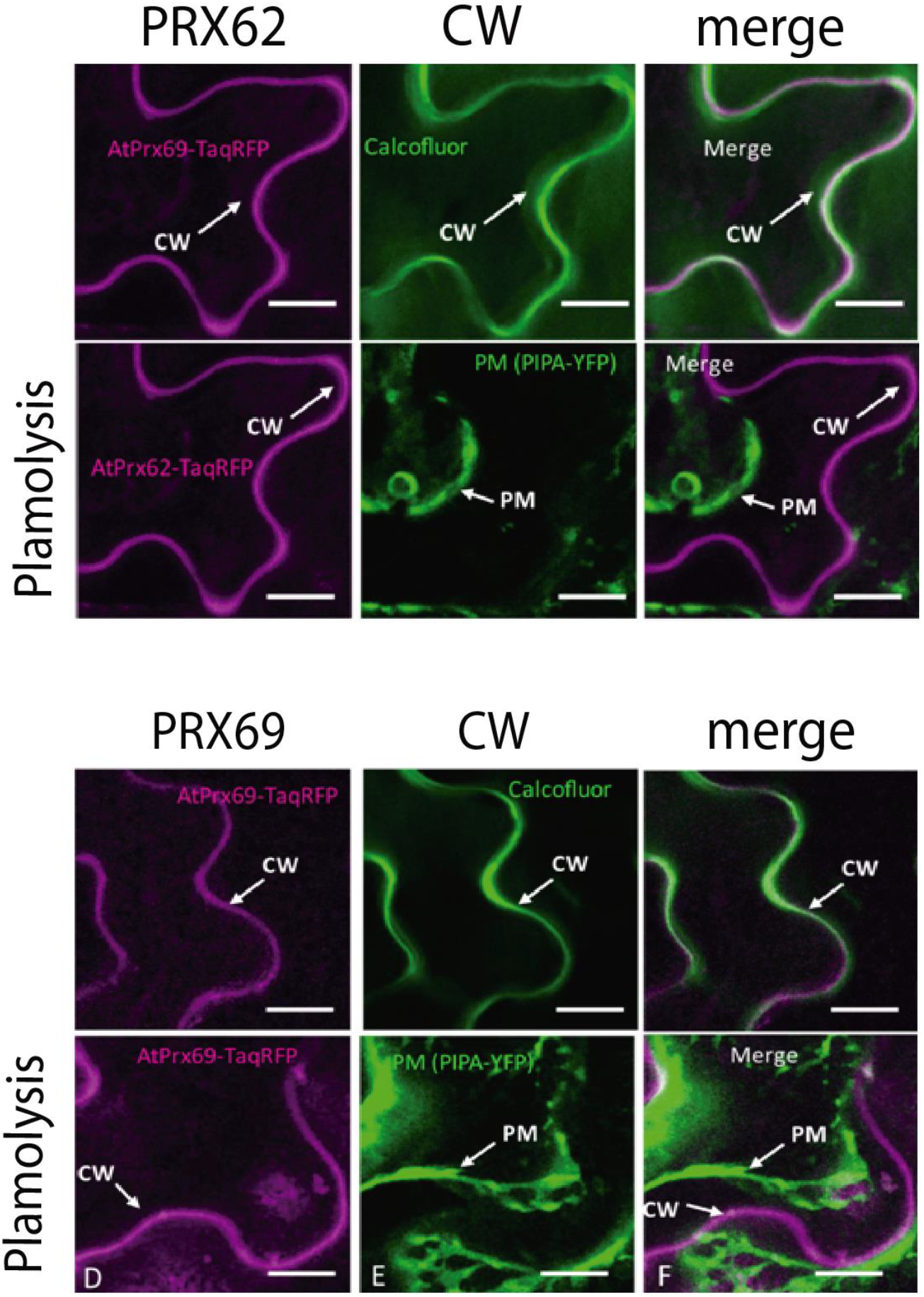
Apoplastic localization of PRX62 and PRX69. Confocal laser scanning fluorescence signals from *Nicotiana benthamiana* plasmolyzed leaf epidermal cells co-expressing 35SproPRX62-TagRFP or 35SproPRX69-TagRFP (magenta channel, left panels) together with the plasma membrane marker Yellow fluorescent protein (YFP)-tagged Plasma Membrane Aquaporin (PIP2A-YFP) (green channel, central panels). Both channels merged (white signal, right panels). The top line corresponds to a single confocal section whereas the bottom line corresponds to the maximum intensity z projection of six confocal sections. Scale bars= 50 μm. CW= cell wall; PM = plasma membrane.

**Figure S6.**
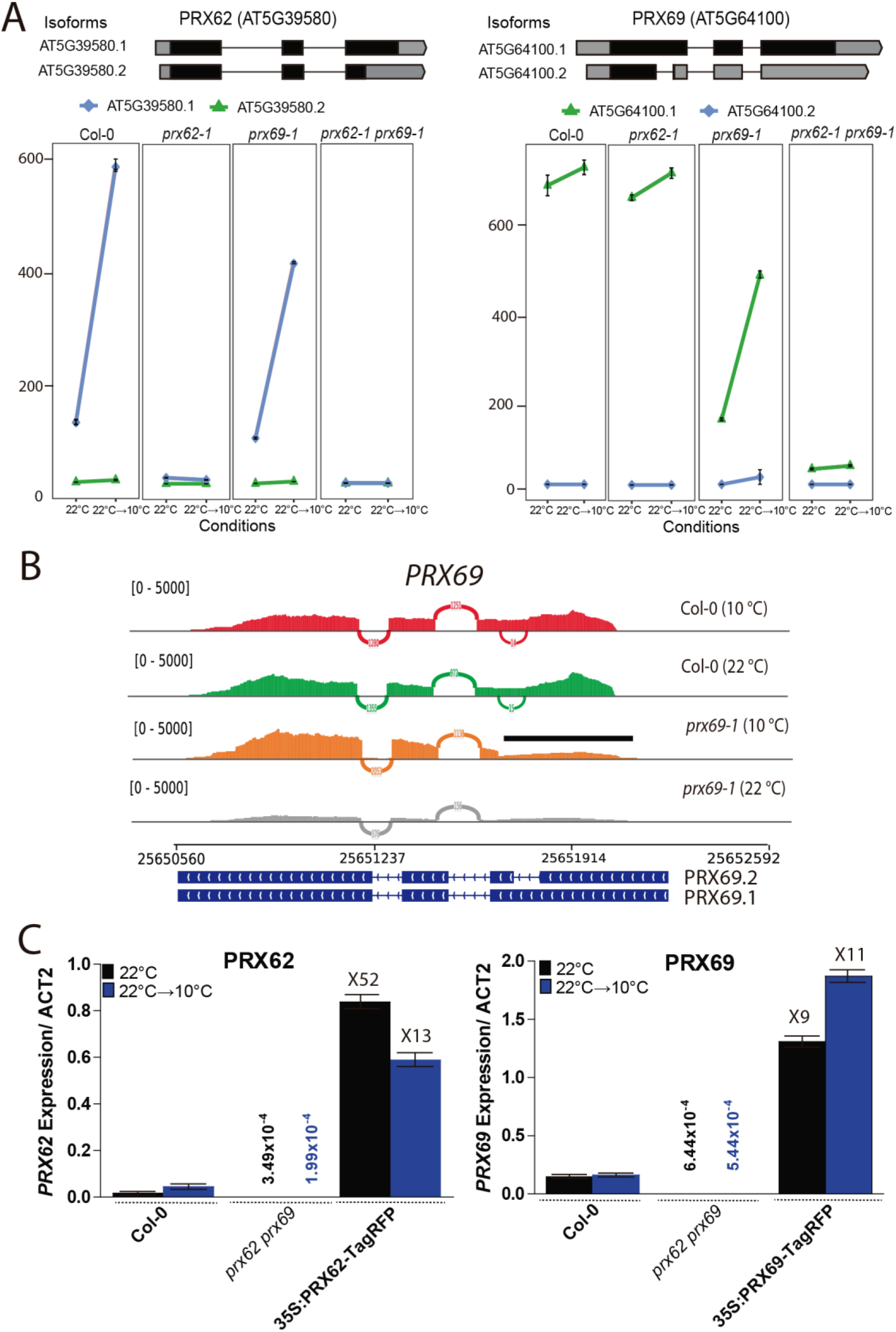
Expression analysis by RNA-seq and Real Time PCR of *PRX62* and *PRX69*. (A) Isoforms expression of *PRX62* and *PRX69* in Col-0, *prx62-1*, *prx69-1* and *prx62-1 prx69-1* double mutant determined by RNA-seq. Reads in *PRX69* gene in *prx69-1* mutant backgrounds showed a truncated version being expressed. Isoforms’ schemes were adapted from boxify (https://boxify.boku.ac.at/). TPM = Transcripts Per Kilobase Million. (B) Sashimi plots of PRX69 indicate the expression of a truncated RNA in the *prx69-1* mutant. Sashimi plots (adapted from IGV) show the coverage for each alignment track (Col-0 and the *prx69-1* mutant) plotted as a histogram; arcs represent splice junctions connecting exons. Alternative splicing isoforms for PRX69 are displayed below. The line on top of the graph highlights the region of the RNA that shows low coverage or low expression. (C) Levels of *PRX62* and *PRX69* expression in Col-0 roots, *prx62-1 prx69-1* double mutant and over-expressor *PRX62* and *PRX69* lines. *ACT2* was use as a housekeeping gene. Three biological replicates and three technical replicates per experiment were performed.

**Figure S7.**
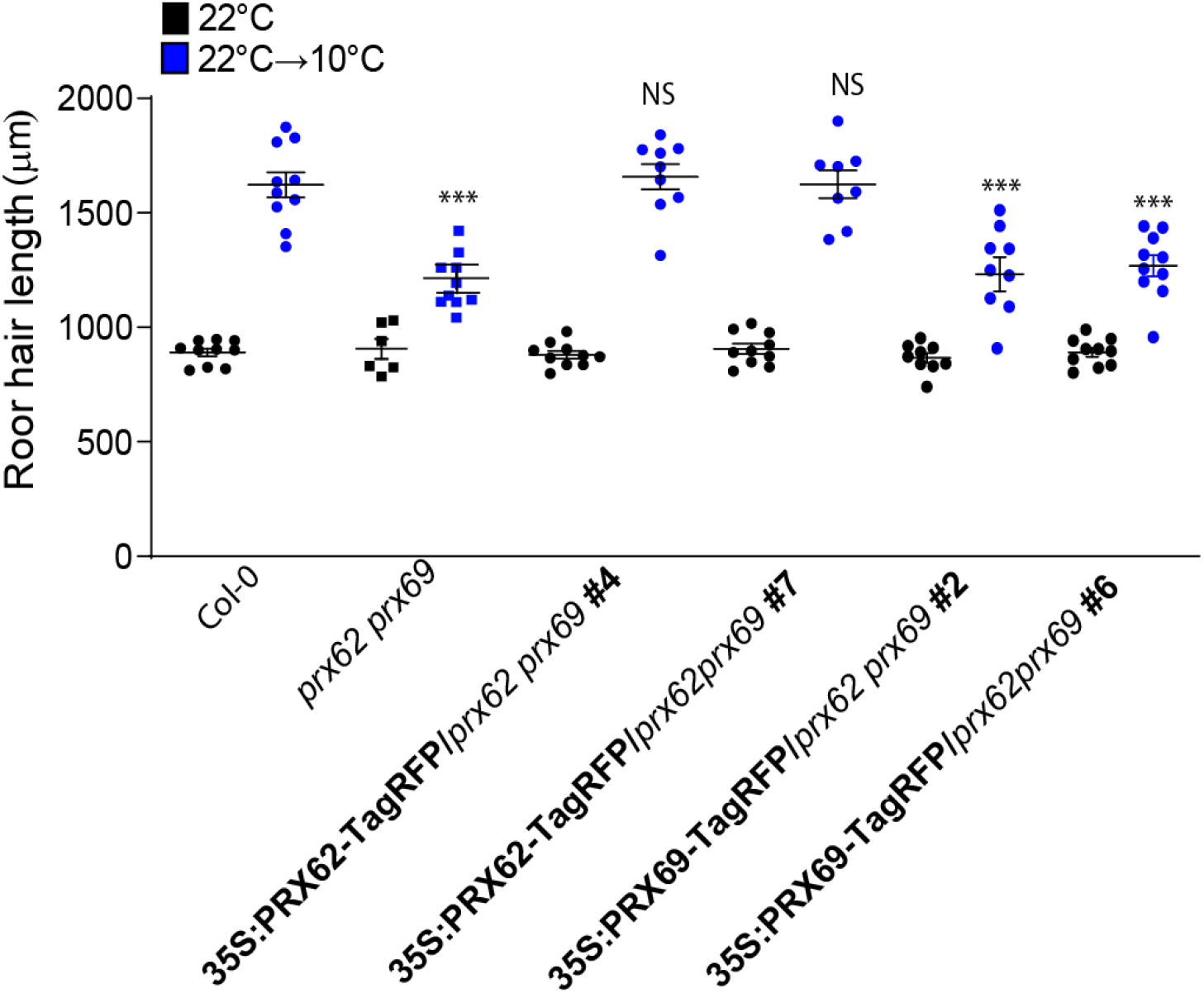
Phenotypic rescue of *prx62 prx69* by overexpression of PRX62 or PRX69. 35SproPRX62 is able to rescue RH growth of *prx62-1 prx69-1* double mutant while 35SproPRX69 failed to rescue of *prx62-1 prx69-1* double mutant. RH length values are the mean of three replicates ± SD. P-value of one-way ANOVA, (***) P<0.001. NS= no significant differences.

**Figure S8.**
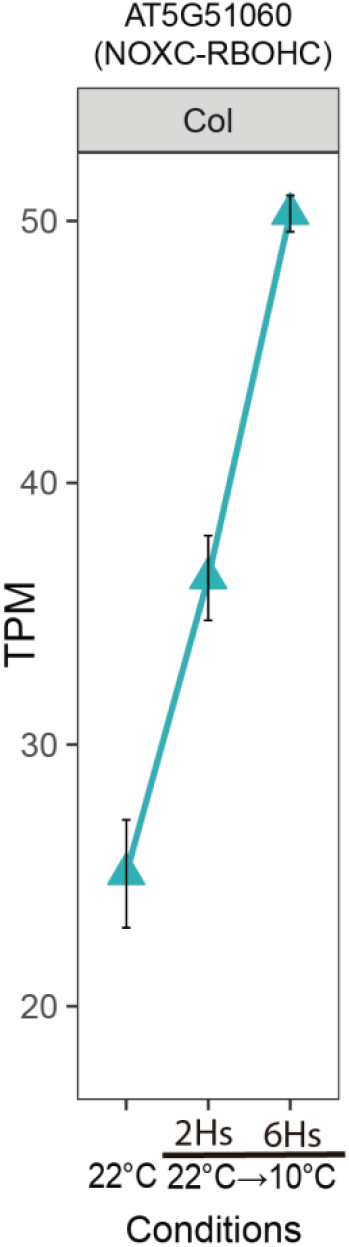
Expression of *NOXC* (*RBOHC/RHD2*) under low-temperature assessed by RNA-seq. TPM = Transcripts Per Kilobase Million.

**Figure S9.**
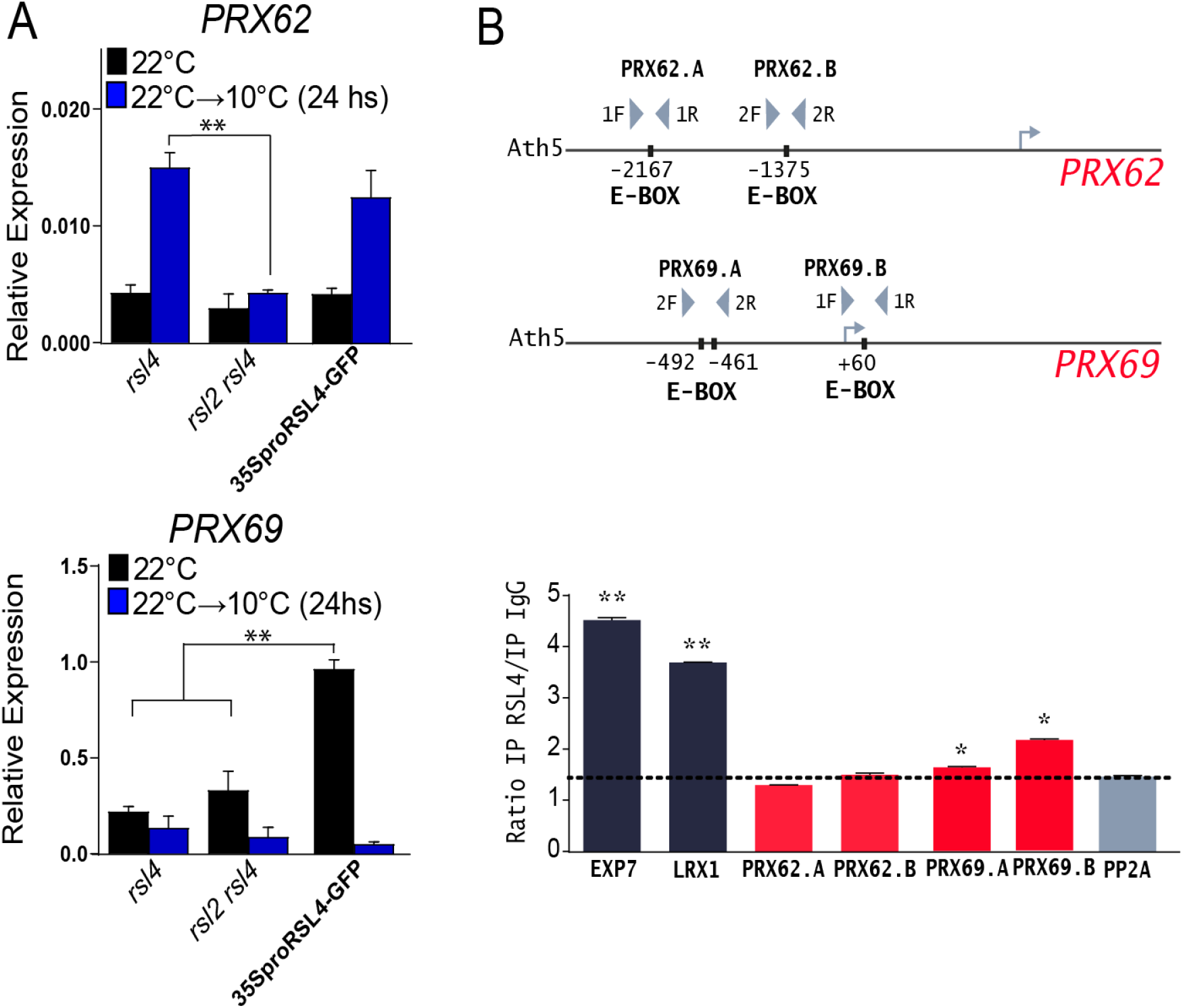
The transcription factor RSL4 controls the expression of *PRX69*. (A) Expression analysis of *PRX62* and *PRX69* by qPCR in Col-0, 35S*_pro_*RSL4, and in the mutants *rsl4*, and *rsl2rsl4* grown at 22°C and then transferred for 24hs at low-temperature treatment (10°C). Three biological replicates and three technical replicates per experiment were performed. P-value of one-way ANOVA,(**) P<0.01. (B) ChIP analysis showing RSL4 binding to RHE-boxes elements (E-boxes) on the *PRX62* and *PRX69* promoter regions. The *PRX62* and *PRX69* promoter regions (pPRXs), the relative positions of RHE-boxes (E-boxes) and ChIP-PCR regions are indicated. The enrichment fold of RSL4-GFP in ChIP-PCR is shown for each region as a ratio with IgG IP taken as a negative control. The enrichment of each region was compared to *PP2A* taken as a negative control and determining the background, indicated as a dotted line. *EXP7* and *LRX1* were taken as positive controls according to Hwang et al. (2017). Significant differences are indicated when appropriate (**P < 0.001, *P < 0.01; t test). The experiment was performed three independent times showing the same trend. The graph represents two technical replicates of one of the independent experiments. F= forward and R=reverse primers.

**Table S2.** PRX62 and PRX69 are transcriptionally induced under low temperature.

| gene name | Log2 (WT 24h/WT 0h) | Gene FDR | Tissue expression |
| --- | --- | --- | --- |
| *PRX05* | 4,68 | 4,7E-21 | endodermis-vascular |
| *PRX04* | 3,31 | 1,5E-43 | endodermis-vascular |
| *PRX37* | 2,10 | 3,2E-15 | endodermis-vascular |
| ***PRX62*** | 2,04 | 6,2E-22 | **RH** |
| *PRX14* | 1,86 | 1,1E-05 | endodermis-vascular |
| ***PRX69*** | 1,76 | 1,4E-35 | **RH** |

FDR= False Discovery Rate. Extracted from Schlaen et al. (2015) where seedlings were grown at 22°C for 9 days and then transferred to 10°C for 24 hs. RH = root hair.

**Table S3.**
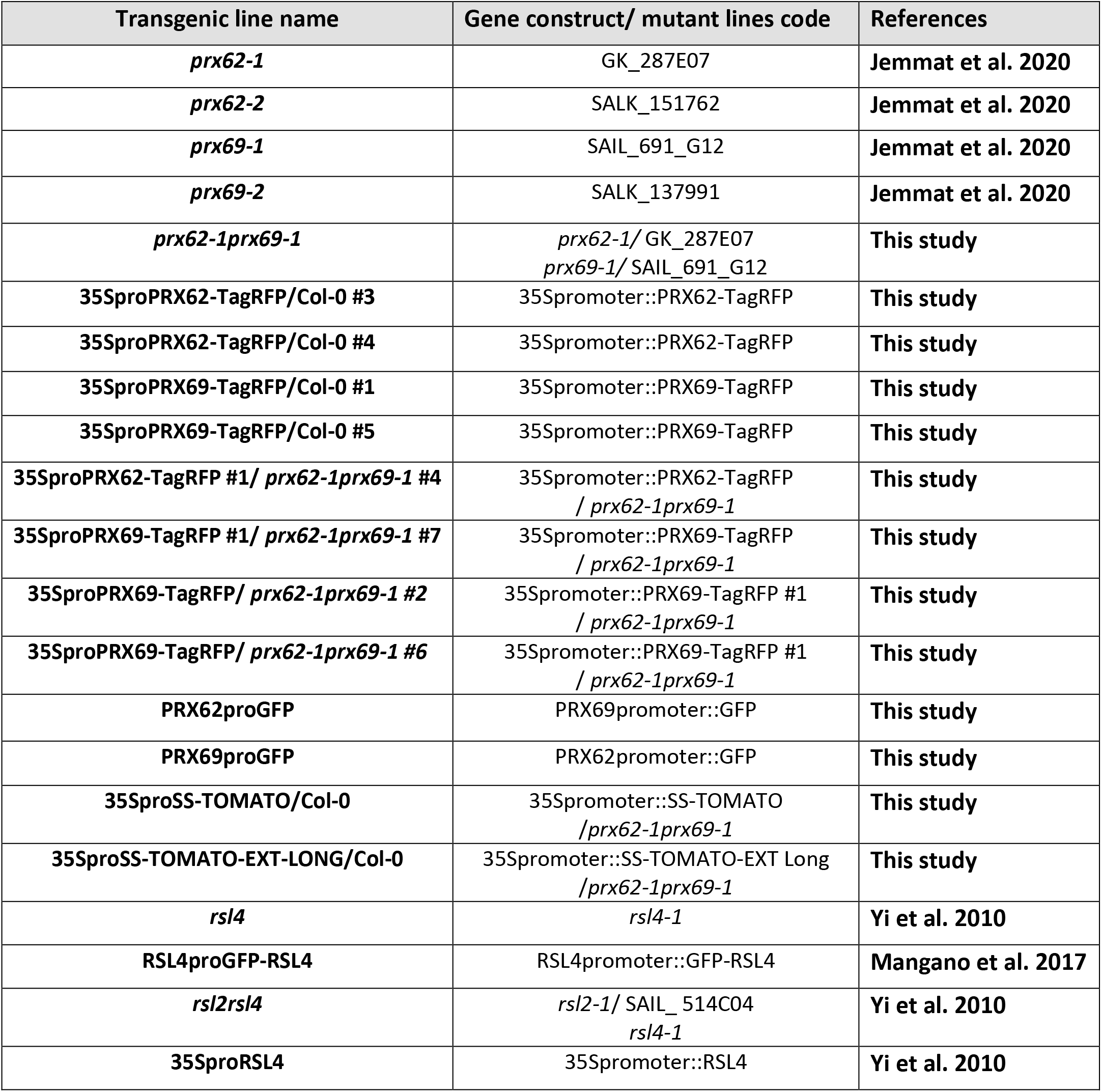
Mutants and transgenic lines generated and used in this study.

**Table S4.**
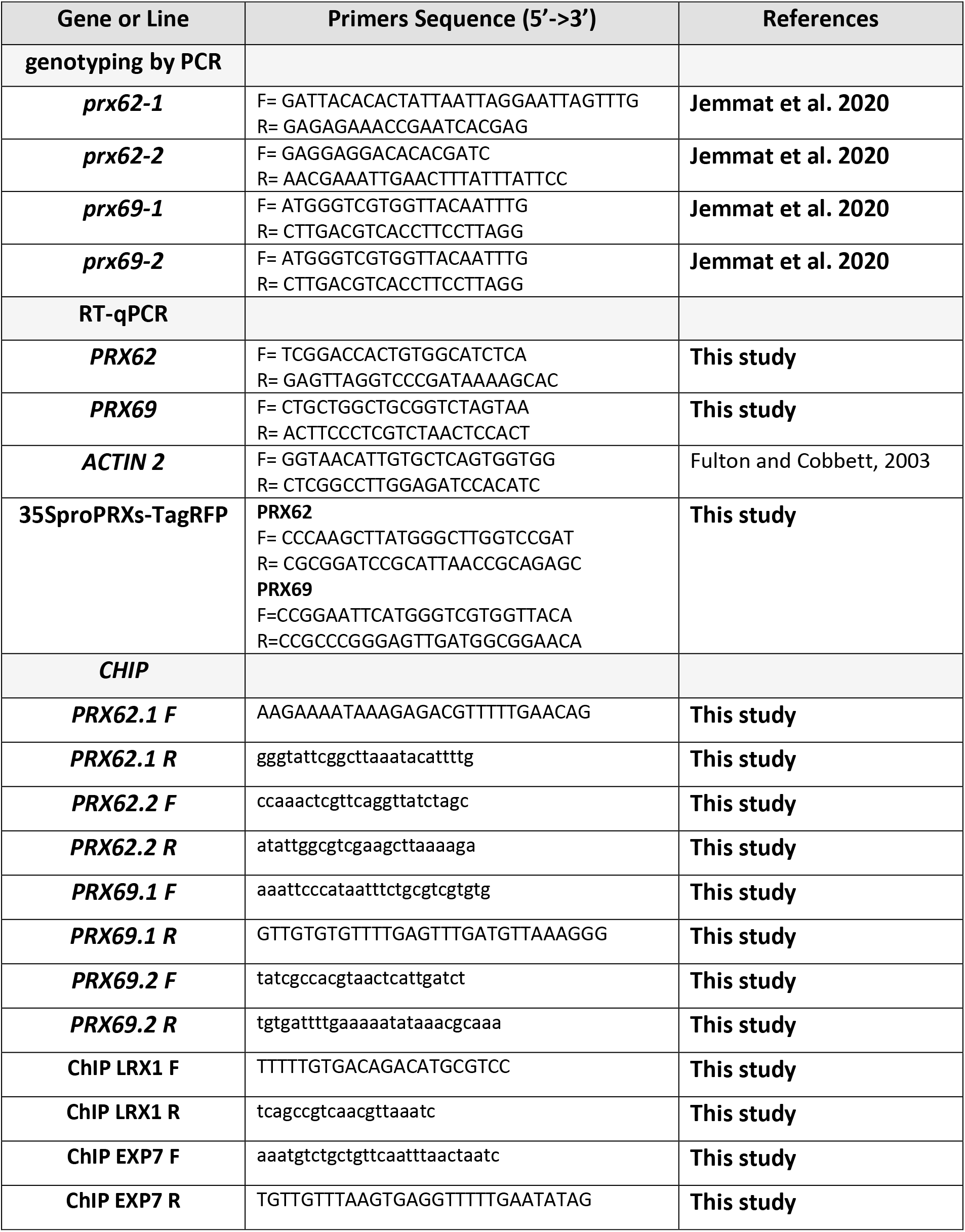
List of Primers used in this study.

## REFERENCES

Ariel, F. et al. R-Loop Mediated trans Action of the APOLO Long Noncoding RNA. Mol. Cell 77, 1055–1065.e4 (2020).

Barah, P. et al. Genome-scale cold stress response regulatory networks in ten Arabidopsis thaliana ecotypes. BMC Genomics 14, 1–16 (2013).

Bashline, L., Lei, L., Li, S. & Gu, Y. Cell wall, cytoskeleton, and cell expansion in higher plants. Mol. Plant 7, 586–600 (2014).

Baumberger, N., Ringli, C. & Keller, B. The chimeric leucine-rich repeat/extensin cell wall protein LRX1 is required for root hair morphogenesis in Arabidopsis thaliana. Genes Dev. 15, 1128–1139 (2001).

Baumberger, N., Steiner, M., Ryser, U., Keller, B. & Ringli, C. Synergistic interaction of the two paralogous Arabidopsis genes LRX1 and LRX2 in cell wall formation during root hair development. Plant J. 35, 71–81 (2003).

Benjamini, Y. & Yekutieli, D. The Control of the False Discovery Rate in Multiple Testing under Dependency. Ann. Stat. 29, 1165–1188 (2001).

Benjamini, Y. & Hochberg, Y. Controlling the False Discovery Rate: A Practical and Powerful Approach to Multiple Testing. J. R. Stat. Soc. Ser. B 57, 289–300 (1995).

Bernards, M. A. et al. Biochemical characterization of the suberization-associated anionic peroxidase of potato. Plant Physiol. 121, 135–46 (1999).

Bhosale, R. et al. A mechanistic framework for auxin dependent Arabidopsis root hair elongation to low external phosphate. Nat. Commun. 9, 1–9 (2018).

Bidhendi, A. J. & Geitmann, A. Relating the mechanics of the primary plant cell wall to morphogenesis. J. Exp. Bot. 67, 449–61 (2016).

Bourbousse, C., Vegesna, N. & Law, J. A. SOG1 activator and MYB3R repressors regulate a complex DNA damage network in Arabidopsis. Proc. Natl. Acad. Sci. U. S. A. 115, E12453–E12462 (2018).

Bullard, J. H., Purdom, E., Hansen, K. D. & Dudoit, S. Evaluation of statistical methods for normalization and differential expression in mRNA-Seq experiments. BMC Bioinformatics 11, 1–13 (2010).

Calixto, C. P. G. et al. Rapid and dynamic alternative splicing impacts the arabidopsis cold response transcriptome[CC-BY]. Plant Cell 30, 1424–1444 (2018).

Cannon, M. C. et al. Self-assembly of the plant cell wall requires an extensin scaffold. Proc. Natl. Acad. Sci. U. S. A. 105, 2226–2231 (2008).

Chapman, J. M., Muhlemann, J. K., Gayomba, S. R. & Muday, G. K. RBOH-Dependent ROS Synthesis and ROS Scavenging by Plant Specialized Metabolites to Modulate Plant Development and Stress Responses. Chemical Research in Toxicology vol. 32 370–396 (2019).

Cosio, C. et al. The class III peroxidase PRX17 is a direct target of the MADS-box transcription factor AGAMOUS-LIKE15 (AGL15) and participates in lignified tissue formation. New Phytol. 213, 250–263 (2017).

Datta, S., Prescott, H. & Dolan, L. Intensity of a pulse of RSL4 transcription factor synthesis determines Arabidopsis root hair cell size. Nat. Plants 1, 1–6 (2015).

Di Rienzo J.A., Casanoves F., Balzarini M.G., Gonzalez L., Tablada M., R. C. W. InfoStat, versión 2008, Grupo InfoStat, FCA, Universidad Nacional de Córdoba, Argentina. (2008).

Ding, Y., Shi, Y. & Yang, S. Advances and challenges in uncovering cold tolerance regulatory mechanisms in plants. New Phytologist vol. 222 1690–1704 (2019).

Ding, Y., Shi, Y. & Yang, S. Molecular Regulation of Plant Responses to Environmental Temperatures. Molecular Plant vol. 13 544–564 (2020).

Dong, W., Kieliszewski, M. & Held, M. A. Identification of the pI 4.6 extensin peroxidase from Lycopersicon esculentum using proteomics and reverse-genomics. Phytochemistry 112, 151–159 (2015).

Dvořák, P., Krasylenko, Y., Zeiner, A., Šamaj, J. & Takáč, T. Signaling Toward Reactive Oxygen Species-Scavenging Enzymes in Plants. Frontiers in Plant Science vol. 11 2178 (2021).

Dunand, C., Crèvecoeur, M. & Penel, C. Distribution of superoxide and hydrogen peroxide in Arabidopsis root and their influence on root development: Possible interaction with peroxidases. New Phytol. 174, 332–341 (2007).

Fabrice, T. N. et al. LRX proteins play a crucial role in pollen grain and pollen tube cell wall development. Plant Physiol. 176, 1981–1992 (2018).

Francoz, E. et al. Pectin Demethylesterification Generates Platforms that Anchor Peroxidases to Remodel Plant Cell Wall Domains. Dev. Cell 48, 261–276.e8 (2019).

Guo, W. et al. 3D RNA-seq: a powerful and flexible tool for rapid and accurate differential expression and alternative splicing analysis of RNA-seq data for biologists. RNA Biol. (2020) doi:10.1080/15476286.2020.1858253.

Hannah, M. A., Heyer, A. G. & Hincha, D. K. A global survey of gene regulation during cold acclimation in Arabidopsis thaliana. PLoS Genet. 1, e26 (2005).

Horton, M. W. et al. Genome-wide patterns of genetic variation in worldwide Arabidopsis thaliana accessions from the RegMap panel. Nat. Genet. 44, 212–216 (2012).

Hossain, M. S. et al. Divergent cytosine DNA methylation patterns in single-cell, soybean root hairs. New Phytol. 214, 808–819 (2017).

Hwang, Y., Choi, H. S., Cho, H. M. & Cho, H. T. Tracheophytes contain conserved orthologs of a basic helix-loop-helix transcription factor that modulate ROOT HAIR SPECIFIC genes. Plant Cell 29, 39–53 (2017).

Jackson, P. A. P. et al. Rapid deposition of extensin during the elicitation of grapevine callus cultures is specifically catalyzed by a 40-kilodalton peroxidase. Plant Physiol. 127, 1065–1076 (2001).

Jacobowitz, J. R., Doyle, W. C. & Weng, J.-K. PRX9 and PRX40 Are Extensin Peroxidases Essential for Maintaining Tapetum and Microspore Cell Wall Integrity during Arabidopsis Anther Development. Plant Cell 31, 848–861 (2019).

Jemmat, A. M. et al. Coordination of five class III peroxidase-encoding genes for early germination events of Arabidopsis thaliana. Plant Sci. 298, 110565 (2020).

Kang, H. M. et al. Efficient Control of Population Structure in Model Organism Association Mapping. Genetics 178, 1709–1723 (2008).

Kang, H. M. et al. Variance component model to account for sample structure in genome-wide association studies. Nat. Genet. 42, 348–354 (2010).

Karimi, M., Inzé, D. & Depicker, A. GATEWAY^TM^ vectors for Agrobacterium-mediated plant transformation. Trends in Plant Science vol. 7 193–195 (2002).

Kim, B. H., Kim, S. Y. & Nam, K. H. Genes encoding plant-specific class III peroxidases are responsible for increased cold tolerance of the brassinosteroid-insensitive 1 mutant. Mol. Cells 34, 539–548 (2012).

Kwon, T. et al. Transcriptional response of Arabidopsis seedlings during spaceflight reveals peroxidase and cell wall remodeling genes associated with root hair development. Am. J. Bot. 102, 21–35 (2015).

Law, C. W., Chen, Y., Shi, W. & Smyth, G. K. Voom: Precision weights unlock linear model analysis tools for RNA-seq read counts. Genome Biol. 15, 1–17 (2014).

Lee, R. D. W. & Cho, H.-T. Auxin, the organizer of the hormonal/environmental signals for root hair growth. Front. Plant Sci. 4, (2013). Mangano, S. et al. Molecular link between auxin and ROS-mediated polar growth. Proc. Natl. Acad. Sci. U. S. A. 114, 5289–5294 (2017).

Lee, Y., Rubio, M. C., Alassimone, J. & Geldner, N. A mechanism for localized lignin deposition in the endodermis. Cell 153, 402–412 (2013).

Lee, Y., Yoon, T.H., Lee, J., Jeon, S.Y., Lee, J.H., Lee, M.K., Chen, H.Z., Yun, J., Oh, S.Y., Wen, X.H., et al. (2018). A lignin molecular brace controls precision processing of cell walls critical for surface integrity in Arabidopsis. Cell 173, 1468–1480.e9.

Leuendorf, J. E., Frank, M. & Schmülling, T. Acclimation, priming and memory in the response of Arabidopsis thaliana seedlings to cold stress. Sci. Rep. 10, 689 (2020).

Mangano, S., Denita-Juarez, S. P., Marzol, E., Borassi, C. & Estevez, J. M. High Auxin and High Phosphate Impact on RSL2 Expression and ROS-Homeostasis Linked to Root Hair Growth in Arabidopsis thaliana. Front. Plant Sci. 9, 1–8 (2018).

Maruyama, K. et al. Identification of cold-inducible downstream genes of the Arabidopsis DREB1A/CBF3 transcriptional factor using two microarray systems. Plant J. 38, 982–993 (2004).

Marzol, E. et al. PRX01, PRX44, and PRX73 are class-III extensin-related peroxidases that modulates root hair growth in Arabidopsis thaliana. bioRxiv 2020.02.04.932376 (2020) doi:10.1101/2020.02.04.932376.

Marzol, E. et al. Filling the Gaps to Solve the Extensin Puzzle. Mol. Plant 11, 645–658 (2018). Mol Plant. 11(5):645–658.

Moison, M. et al. The lncRNA APOLO interacts with the transcription factor WRKY42 to trigger root hair cell expansion in response to cold. Mol. Plant 0, (2021).

Nakagawa, T. et al. Development of series of gateway binary vectors, pGWBs, for realizing efficient construction of fusion genes for plant transformation. J. Biosci. Bioeng. 104, 34–41 (2007).

Nurhasanah Ritonga, F. & Chen, S. Physiological and molecular mechanism involved in cold stress tolerance in plants. Plants vol. 9 560 (2020).

Orman-Ligeza, B. et al. RBOH-mediated ROS production facilitates lateral root emergence in Arabidopsis. Development 143, 3328–39 (2016).

Pacheco, J. M. et al. The lncRNA APOLO and the transcription factor WRKY42 target common cell wall EXTENSIN encoding genes to trigger root hair cell elongation. Plant Signal. Behav. 16, 1920191 (2021).

Passardi, F., Longet, D., Penel, C. & Dunand, C. The class III peroxidase multigenic family in rice and its evolution in land plants. Phytochemistry 65, 1879–1893 (2004).

Passardi, F., Penel, C. & Dunand, C. Performing the paradoxical: How plant peroxidases modify the cell wall. Trends in Plant Science vol. 9 534–540 (2004).

Patro, R., Duggal, G., Love, M. I., Irizarry, R. A. & Kingsford, C. Salmon provides fast and bias-aware quantification of transcript expression. Nat. Methods 14, 417–419 (2017).

Penfield, S. Temperature perception and signal transduction in plants. New Phytologist vol. 179 615–628 (2008).

Quint, M. et al. Molecular and genetic control of plant thermomorphogenesis. Nature Plants vol. 2 (2016).

Pereira, C. S. et al. Extensin network formation in Vitis vinifera callus cells is an essential and causal event in rapid and H2O2-induced reduction in primary cell wall hydration. BMC Plant Biol. 11, 106 (2011).

Pfaffl, M. W. A new mathematical model for relative quantification in real-time RT-PCR. Nucleic Acids Res. 29, 45e – 45 (2001).

Pfaffl, M. W., Horgan, G. W. & Dempfle, L. Relative expression software tool (REST) for group-wise comparison and statistical analysis of relative expression results in real-time PCR. Nucleic Acids Res. 30, 36e – 36 (2002).

Price, N. J. et al. A Biochemical and Molecular Characterization of LEP1, an Extensin Peroxidase from Lupin. J. Biol. Chem. 278, 41389–41399 (2003).

Ringli, C. The hydroxyproline-rich glycoprotein domain of the Arabidopsis LRX1 requires Tyr for function but not for insolubilization in the cell wall. Plant J. 63, 662–669 (2010).

Ritchie, M. E. et al. Limma powers differential expression analyses for RNA-sequencing and microarray studies. Nucleic Acids Res. 43, e47 (2015).

Saraçli, S., Doǧan, N. & Doǧan, I. Comparison of hierarchical cluster analysis methods by cophenetic correlation. J. Inequalities Appl. 2013, 1–8 (2013).

Schlaen, R. G. et al. The spliceosome assembly factor GEMIN2 attenuates the effects of temperature on alternative splicing and circadian rhythms. Proc. Natl. Acad. Sci. U. S. A. 112, 9382–9387 (2015).

Schnabelrauch, L. S., Kieliszewski, M., Upham, B. L., Alizedeh, H. & Lamport, D. T. A. Isolation of pl 4.6 extensin peroxidase from tomato cell suspension cultures and identification of Val-Tyr-Lys as putative intermolecular cross-link site. Plant J. 9, 477–489 (1996).

Sede, A. R. et al. Arabidopsis pollen extensins LRX are required for cell wall integrity during pollen tube growth. FEBS Lett. 592, 233–243 (2018).

Seren, Ü. et al. GWAPP: A Web Application for Genome-Wide Association Mapping in Arabidopsis. Plant Cell 24, 4793–4805 (2012).

Shin, J. H., Blay, S., McNeney, B. & Graham, J. LDheatmap: An R function for graphical display of pairwise linkage disequilibria between single nucleotide polymorphisms. J. Stat. Softw. 16, 1–9 (2006).

Soneson, C., Love, M. I. & Robinson, M. D. Differential analyses for RNA-seq: Transcript-level estimates improve gene-level inferences. F1000Research 4, 1521 (2016).

Srivastava, L. M. Cell Wall, Cell Division, and Cell Growth. in Plant Growth and Development 23–74 (Elsevier, 2002). doi:10.1016/b978-012660570-9/50142-8.

Turner, S. Qqman: Q-Q and Manhattan Plots for GWAS Data. R Package Version 0.1.8. 5 https://cran.r-project.org/package=qqman (2021).

Valdés-López, O. et al. Soybean roots grown under heat stress show global changes in their transcriptional and proteomic profiles. Front. Plant Sci. 7, 517 (2016).

Valério, L., De Meyer, M., Penel, C. & Dunand, C. Expression analysis of the Arabidopsis peroxidase multigenic family. Phytochemistry 65, 1331–1342 (2004).

Velasquez, S. M. et al. Low Sugar Is Not Always Good: Impact of Specific O-Glycan Defects on Tip Growth in Arabidopsis. Plant Physiol. 168, 808–813 (2015).

Velasquez, S. M. et al. O-glycosylated cell wall proteins are essential in root hair growth. Science (80- . ). 332, 1401–1403 (2011).

Waese, J. et al. ePlant: Visualizing and exploring multiple levels of data for hypothesis generation in plant biology. Plant Cell 29, 1806–1821 (2017).

Wang, X. et al. Pollen-expressed leucine-rich repeat extensins are essential for pollen germination and growth. Plant Physiol. 176, 1993–2006 (2018).

Wojtaszek, P., Trethowan, J. & Bolwell, G. P. Reconstitution in vitro of the components and conditions required for the oxidative cross-linking of extracellular proteins in French bean (Phaseolus vulgaris L.). FEBS Lett. 405, 95–98 (1997).

Yaqoob, A. et al. Dual functions of Expansin in cell wall extension and compression during cotton fiber development. Biologia (Bratisl). 75, 2093–2101 (2020).

Yi, K., Menand, B., Bell, E. & Dolan, L. A basic helix-loop-helix transcription factor controls cell growth and size in root hairs. Nat. Genet. 42, 264–267 (2010).

Zhang, S. et al. Multiple phytohormones promote root hair elongation by regulating a similar set of genes in the root epidermis in Arabidopsis. J. Exp. Bot. 67, 6363–6372 (2016).

Zhang, Z. et al. Mixed linear model approach adapted for genome-wide association studies. Nat. Genet. 42, 355–360 (2010).

Zhang, R. et al. A high quality Arabidopsis transcriptome for accurate transcript-level analysis of alternative splicing. Nucleic Acids Res. 45, 5061–5073 (2017).

