## Supplementary material for "Apoplastic class III peroxidases PRX62 and PRX69 regulate ROS-homeostasis and cell wall associated extensins linked to root hair growth at low-temperature in *Arabidopsis thaliana*": Table S1

| Stock_N | Genotype_ID | Name | RH length 10C µm | SD | N=Plants | RH length 22C µm | SD | N=Plants |
|---|---|---|---|---|---|---|---|---|
| 650 | 7404 | Wc-1 | 2218,1 | 46,6 | 5 | 542,33 | 23,74 | 6 |
| 659 | 7328 | Sf-2 | 2061,6 | 384,8 | 12 | 639,53 | 188,17 | 16 |
| 651 | 6964 | Spr1-2 | 1911,3 | 252,2 | 15 | 838,35 | 80,84 | 12 |
| 671 | 6957 | Pu2-8 | 1758,4 | 376,4 | 13 | 907,94 | 172,17 | 6 |
| 664 | 7242 | Lo-2 | 1741,1 | 150,9 | 6 | 663,77 | 114,04 | 6 |
| 535 | 7033 | Buckhorn Pass | 1666,9 | 339,6 | 6 | 535,97 | 114,62 | 7 |
| 629 | 7121 | En-T | 1586,4 | 156,0 | 7 | 747,08 | 124,38 | 12 |
| 619 | 8259 | Bå5-1 | 1476,2 | 73,2 | 5 | 585,51 | 220,98 | 9 |
| 657 | 7149 | Gu-0 | 1412,0 | 167,2 | 5 | 715,52 | 77,54 | 7 |
| 423 | 8271 | Bu-0 | 1395,5 | 252,6 | 7 | 748,60 | 121,79 | 9 |
| 531 | 7504 | PHW-33 | 1378,3 | 188,8 | 6 | 511,33 | 59,45 | 9 |
| 542 | 7461 | H55 | 1229,8 | 281,0 | 5 | 496,08 | 91,30 | 5 |
| 132 | 6923 | HR-10 | 1169,6 | 89,8 | 6 | 416,52 | 159,22 | 9 |
| 672 | 7519 | Ömö2-3 | 1166,1 | 151,3 | 5 | 1028,27 | 102,69 | 5 |
| 462 | 8296 | Gd-1 | 1127,0 | 236,7 | 7 | 781,62 | 76,78 | 9 |
| 613 | 8243 | PHW-2 | 1118,3 | 176,9 | 14 | 754,40 | 162,41 | 21 |
| 214 | 7408 | Wt-3 | 1075,0 | 221,1 | 11 | 496,90 | 124,86 | 19 |
| 603 | 8334 | Lu-1 | 1067,8 | 133,7 | 8 | 260,22 | 62,14 | 9 |
| 627 | 8270 | Bs-1 | 1063,5 | 225,1 | 7 | 853,17 | 72,23 | 7 |
| 79 | 6910 | Ct-1 | 1036,2 | 123,1 | 18 | 496,48 | 122,90 | 15 |
| 89 | 8387 | St-0 | 1035,6 | 196,7 | 16 | 532,69 | 190,09 | 15 |
| 221 | 6924 | HR-5 | 1011,9 | 135,7 | 6 | 508,92 | 212,67 | 13 |
| 609 | 8423 | Hov2-1 | 1010,8 | 421,6 | 6 | 494,00 | 111,72 | 9 |
| 197 | 6943 | NFA-10 | 977,2 | 173,8 | 16 | 819,23 | 76,31 | 17 |
| 82 | 362 | TOU-C-3 | 969,8 | 115,5 | 17 | 515,44 | 249,96 | 20 |
| 224 | 6926 | Kin-0 | 965,6 | 77,2 | 9 | 512,63 | 209,43 | 13 |
| 611 | 7492 | PHW-22 | 965,1 | 203,9 | 12 | 390,57 | 118,54 | 12 |
| 637 | 7292 | Pa-3 | 961,9 | 157,1 | 7 | 403,27 | 76,86 | 7 |
| 220 | 8357 | Pla-0 | 951,7 | 112,3 | 13 | 388,93 | 51,42 | 18 |
| 644 | 7080 | Co-4 | 938,1 | 153,7 | 6 | 578,94 | 130,90 | 8 |
| 646 | 8426 | Ull1-1 | 916,6 | 277,6 | 7 | 647,44 | 113,04 | 7 |
| 473 | 7256 | Mh-1 | 901,2 | 341,0 | 13 | 616,26 | 96,22 | 13 |
| 530 | 7109 | Ema-1 | 893,3 | 195,9 | 12 | 435,06 | 74,74 | 14 |
| 146 | 5381 | UKNW06-060 | 891,7 | 189,2 | 8 | 371,35 | 78,56 | 5 |
| 16 | 5264 | UKSE06-520 | 881,8 | 310,0 | 8 | 556,36 | 144,78 | 7 |
| 279 | 6960 | Ren-11 | 875,7 | 314,0 | 8 | 435,36 | 146,33 | 7 |
| 522 | 98021 | Bozen-1 | 857,2 | 218,3 | 13 | 417,02 | 78,06 | 13 |
| 133 | 6909 | Col-0 | 847,9 | 197,2 | 30 | 373,84 | 95,48 | 20 |
| 2 | 7135 | Fr-4 | 847,8 | 467,4 | 5 | 302,99 | 88,52 | 10 |
| 515 | 6944 | NFA-8 | 840,9 | 214,3 | 15 | 414,52 | 75,87 | 5 |
| 615 | 98053 | Mer-6 | 839,5 | 179,7 | 16 | 446,06 | 209,77 | 15 |
| 149 | 5565 | UKNW06-386 | 833,1 | 52,5 | 5 | 433,80 | 125,58 | 4 |
| 605 | 8395 | Tu-0 | 830,8 | 86,8 | 5 | 797,63 | 167,95 | 7 |
| 549 | 6911 | Cvi-0 | 830,4 | 119,6 | 5 | 660,64 | 186,61 | 10 |
| 225 | 7265 | Nd-0 | 823,0 | 307,6 | 15 | 234,10 | 112,70 | 19 |
| 218 | 8610 | 11ME1.32 | 819,9 | 178,5 | 12 | 424,02 | 49,59 | 14 |

| | | | | | | | | |
|---|---|---|---|---|---|---|---|---|
| 628 | 6994 | Ann-1 | 815,8 | 242,1 | 22 | 610,05 | 117,20 | 24 |
| 543 | 7079 | Co-3 | 803,4 | 144,8 | 15 | 569,22 | 88,06 | 9 |
| 447 | 8312 | Is-0 | 789,9 | 146,4 | 5 | 436,03 | 78,31 | 5 |
| 607 | 6449 | ZdrI2-25 | 780,6 | 234,6 | 5 | 508,10 | 56,27 | 6 |
| 538 | 8337 | Mir-0 | 780,3 | 212,9 | 9 | 653,61 | 92,67 | 9 |
| 617 | 8290 | En-1 | 778,5 | 259,1 | 5 | 428,55 | 55,75 | 5 |
| 653 | 7195 | Kl-1 | 773,3 | 214,6 | 6 | 458,76 | 82,38 | 7 |
| 623 | 6931 | Kz-9 | 770,6 | 159,9 | 5 | 292,64 | 63,57 | 3 |
| 658 | 7228 | Li5:3 | 758,7 | 186,8 | 6 | 590,06 | 99,76 | 8 |
| 227 | 7276 | Ob-0 | 754,0 | 244,4 | 14 | 706,78 | 139,79 | 15 |
| 642 | 8215 | Fei-0 | 750,5 | 132,4 | 5 | 435,84 | 122,03 | 6 |
| 655 | 8258 | Bå4-1 | 741,7 | 222,3 | 12 | 504,44 | 68,96 | 21 |
| 98 | 8249 | Vimmerby | 738,1 | 294,6 | 15 | 557,44 | 61,93 | 12 |
| 199 | 7239 | Ll-2 | 733,1 | 135,1 | 15 | 408,36 | 47,61 | 5 |
| 393 | 7110 | Ede-1 | 719,2 | 272,9 | 5 | 549,81 | 13,50 | 2 |
| 540 | 6936 | Lz-0 | 715,6 | 209,9 | 14 | 232,14 | 50,10 | 6 |
| 135 | 6966 | Sq-1 | 676,3 | 82,5 | 12 | 549,15 | 58,92 | 12 |
| 256 | 6039 | Hovdala-2 | 675,6 | 68,3 | 11 | 568,70 | 83,56 | 13 |
| 157 | 6979 | Wei-0 | 670,8 | 212,2 | 7 | 346,00 | 50,09 | 9 |
| 124 | 6985 | Zdr-6 | 651,1 | 107,0 | 14 | 526,86 | 232,62 | 18 |
| 248 | 7165 | Hn-0 | 648,3 | 159,1 | 5 | 273,82 | 56,83 | 5 |
| 521 | 6448 | ZdrI2-24 | 641,7 | 117,0 | 5 | 406,30 | 55,95 | 6 |
| 222 | 8240 | Kulturen-1 | 632,2 | 66,4 | 5 | 486,47 | 114,16 | 7 |
| 622 | 8254 | Ang-0 | 626,4 | 120,1 | 11 | 510,68 | 96,56 | 8 |
| 139 | 6968 | Tamm-2 | 626,1 | 151,1 | 18 | 326,41 | 85,14 | 19 |
| 30 | 98012 | Valsi-1 | 615,7 | 156,9 | 17 | 327,26 | 137,82 | 13 |
| 247 | 6939 | Mt-0 | 613,9 | 154,3 | 5 | 241,58 | 70,83 | 6 |
| 119 | 7518 | Ömö2-1 | 612,9 | 106,1 | 9 | 367,36 | 36,87 | 9 |
| 122 | 6973 | Ull 2-3 | 602,8 | 177,4 | 17 | 264,04 | 57,35 | 9 |
| 210 | 6977 | Van-0 | 597,5 | 59,4 | 16 | 358,40 | 89,72 | 17 |
| 209 | 7355 | Tiv-1 | 589,2 | 136,4 | 12 | 305,72 | 55,89 | 10 |
| 624 | 8388 | Stw-0 | 553,6 | 174,4 | 17 | 242,42 | 66,67 | 18 |
| 523 | 6933 | LL-0 | 537,3 | 141,3 | 5 | 324,51 | 46,88 | 4 |
| 91 | 6906 | C24 | 533,5 | 231,1 | 15 | 448,25 | 119,98 | 16 |
| 24 | 98024 | Bolin-1 | 525,2 | 100,2 | 7 | 270,44 | 60,29 | 8 |
| 643 | 6929 | Kondara | 507,0 | 141,6 | 13 | 517,30 | 124,26 | 12 |
| 237 | 6946 | Oy-0 | 502,9 | 131,6 | 6 | 288,76 | 111,75 | 7 |
| 231 | 8796 | 11 PNA4.101 | 495,0 | 104,5 | 10 | 310,57 | 38,41 | 11 |
| 159 | 6983 | Yo-0 | 493,4 | 242,0 | 6 | 381,11 | 105,95 | 17 |
| 257 | 7520 | Lp 2-2 | 492,3 | 137,3 | 10 | 256,36 | 46,23 | 10 |
| 21 | 98059 | Sha | 473,6 | 146,4 | 10 | 321,24 | 119,98 | 7 |
| 255 | 6988 | Alc-0 | 449,5 | 103,2 | 9 | 358,05 | 127,51 | 10 |
| 141 | 6930 | Kz-1 | 444,6 | 129,0 | 5 | 327,11 | 66,06 | 4 |
| 63 | 98018 | Sij-2 | 429,8 | 207,2 | 12 | 306,68 | 100,35 | 11 |
| 576 | 6915 | Ei-2 | 422,1 | 117,2 | 10 | 419,39 | 136,10 | 9 |
| 590 | 7017 | Bla-11 | 409,1 | 70,1 | 7 | 393,78 | 70,26 | 7 |
| 19 | 98043 | Krazo-2 | 399,9 | 134,4 | 7 | 355,71 | 100,88 | 10 |

| | | | | | | | | |
|---|---|---|---|---|---|---|---|---|
| 235 | 6962 | Shahdara | 339,1 | 137,0 | 5 | 313,42 | 52,86 | 3 |
| 587 | 7386 | Vi-0 | 304,1 | 84,1 | 7 | 323,27 | 104,19 | 6 |
| 584 | 7403 | Wei-1 | 296,1 | 72,6 | 11 | 306,66 | 69,54 | 10 |
| 20 | 98047 | Angit-1 | 290,8 | 103,7 | 15 | 219,06 | 40,86 | 16 |
| 233 | 5232 | UKSE06-466 | 270,4 | 69,6 | 7 | 207,03 | 52,91 | 8 |
| 592 | 7238 | Ll-1 | 252,6 | 55,3 | 8 | 270,59 | 32,27 | 7 |
| 362 | 5887 | DraIV 1-5 | 233,2 | 177,9 | 5 | 266,17 | 52,59 | 4 |
| 569 | 7024 | Bla-6 | 217,9 | 51,8 | 8 | 224,26 | 41,40 | 5 |
| 54 | 98050 | Ciste-2 | 217,5 | 58,7 | 12 | 214,12 | 52,90 | 13 |
| 12 | 98051 | Lag2.2 | 188,5 | 85,8 | 9 | 172,50 | 42,84 | 11 |
| 552 | 8411 | Rd-0 | 186,3 | 36,8 | 9 | 193,09 | 35,36 | 9 |
| 541 | 7147 | Gie-0 | 172,0 | 42,5 | 5 | 176,80 | 51,74 | 6 |
| 468 | 7102 | Do-0 | 161,6 | 41,9 | 6 | 227,21 | 112,74 | 4 |
| 586 | 7352 | Te-0 | 148,6 | 57,4 | 5 | 126,10 | 2,02 | 2 |
